# Protein2Text: Providing Rich Descriptions for Protein Sequences

**DOI:** 10.1101/2024.12.04.626777

**Authors:** Edo Dotan, Iris Lyubman, Eran Bacharach, Tal Pupko, Yonatan Belinkov

## Abstract

Understanding the functionality of proteins has been a focal point of biological research due to their critical roles in various biological processes. Unraveling protein functions is essential for advancements in medicine, agriculture, and biotechnology, enabling the development of targeted therapies, engineered crops, and novel biomaterials. However, this endeavor is challenging due to the complex nature of proteins, requiring sophisticated experimental designs and extended timelines to uncover their specific functions. Public large language models (LLMs), though proficient in natural language processing, struggle with biological sequences due to the unique and intricate nature of biochemical data. These models often fail to accurately interpret and predict the functional and structural properties of proteins, limiting their utility in bioinformatics. To address this gap, we introduce BetaDescribe, a collection of models designed to generate detailed and rich textual descriptions of proteins, encompassing properties such as function, catalytic activity, involvement in specific metabolic pathways, subcellular localizations, and the presence of particular domains. The trained BetaDescribe model receives protein sequences as input and outputs a textual description of these properties. BetaDescribe’s starting point was the LLAMA2 model, which was trained on trillions of tokens. Next, we trained our model on datasets containing both biological and English text, allowing biological knowledge to be incorporated. We demonstrate the utility of BetaDescribe by providing descriptions for proteins that share little to no sequence similarity to proteins with functional descriptions in public datasets. We also show that BetaDescribe can be harnessed to conduct *in-silico* mutagenesis procedures to identify regions important for protein functionality without needing homologous sequences for the inference. Altogether, BetaDescribe offers a powerful tool to explore protein functionality, augmenting existing approaches such as annotation transfer based on sequence or structure similarity.

## Introduction

Since the discovery of the first protein sequence (Sanger & Thompson, 1953), researchers have been fascinated by understanding the intricate functionality of proteins. Proteins play a vital role in biological processes, serving as catalysts for chemical reactions, signal transmitters, structural building blocks, and much more. Unraveling the specific functions of proteins is crucial for advancing fields like medicine, agriculture, and biotechnology. By comprehensively understanding how proteins operate, scientists can develop targeted therapies for diseases, engineer crops with desirable traits, and design novel biomaterials. Furthermore, elucidating protein functionality is essential for uncovering the complexities of life itself, offering insights into evolution, cellular dynamics, and organismal behavior. Thus, deciphering protein functionality is a cornerstone in the quest to comprehend the intricacies of biological systems.

Unlocking the functionality of proteins poses a formidable challenge due to the intricate and multifaceted nature of these biomolecules. Researchers often embark on complex wet lab experiments, navigating through a labyrinth of variables and interactions. Designing experiments to elucidate protein functionality requires meticulous planning, innovative techniques, and sophisticated instrumentation. From studying protein-protein interactions to deciphering their three-dimensional structures, each aspect demands tailored experimental approaches. Moreover, the dynamic nature of proteins adds another layer of complexity, as their functions can be influenced by environmental factors, post-translational modifications (PTM), and interactions with other molecules. In addition, when facing an unknown protein, typically, researchers need to design and conduct multiple experiments, each targeting a specific hypothesis. Thus, experimental determination of the functionality of a new protein may take a few years. As a result, the functions of most proteins across all domains of life are computationally predicted.

In the last decade, artificial neural networks have emerged as a powerful paradigm for solving complex problems in different fields (LeCun et al., 2015), such as computer vision (Voulodimos et al., 2018), natural language processing (NLP; Young et al., 2018), speech (Nassif et al., 2019), and structural biology (Jumper et al., 2021). Biological sequences, like natural languages, are composed of discrete characters: letters in human languages, nucleotides / ribonucleotides in DNA / RNA sequences, and amino acids in proteins. These characters form the foundation for more complex structures, such as sentences and genes, which ultimately create documents and genomes (Simon et al., 2024; Dotan et al., 2024). However, there are many differences between the two. While human languages follow known grammatical rules, specific morphological structures, and contextual cues, biological sequences are arranged in highly specific and intricate patterns that convey complex biochemical information. Furthermore, in contrast to biological sequences, evaluating texts written in natural languages, like English, is relatively straightforward; a simple read-through by a native speaker can reveal errors, convey meaning, and provide insights into the text’s quality and coherence. Nevertheless, similar properties of protein sequences and English text allow for adapting NLP-based techniques for protein analyses (Hayes et al., 2024; Lin et al., 2023; Nijkamp et al., 2023; Yuan et al., 2024; Zhang et al., 2024).

In this work, we developed BetaDescribe, a collection of models trained to generate rich textual descriptions of proteins. Given a protein sequence as input, our trained model describes several properties that may include the protein’s function, catalytic activity, subcellular localization, and the PTMs it can undergo. The starting model of BetaDescribe is a LLAMA2 model (Touvron et al., 2023), trained on trillions of English tokens (and thus, lacking expert biological knowledge). We further trained this model on more than 120 billion tokens containing protein knowledge extracted from UniProt. We tested the performance of BetaDescribe by evaluating the differences between its generated descriptions for protein sequences with a known function and the functions reported in UniProt. We also demonstrate its applicability for predicting the function of proteins with insignificant sequence similarity to any of the proteins used for training. BetaDescribe acts as a bridge between protein sequences and their rich functional descriptions.

## New Approaches

### Outline

BetaDescribe is a collection of deep-learning models designed to generate and validate optimal descriptions of proteins. This collection comprises three components: ’*generator*’, ’*validators*’, and ’*judge*’ (Figure 1). The generator creates rich and detailed candidate textual descriptions for each protein. The validators predict a few simple properties of proteins (e.g., the subcellular localization of the protein). The judge receives a candidate description (from the generator) and the predicted properties (from the validators) and rejects or accepts the candidate. The generator and the validators process the protein sequence and generate textual (English) descriptions and properties, respectively. The judge processes English text only. Specifically, we trained a large model (7 billion parameters) as the generator, and smaller models (150 million parameters) as validators (since the generation of text is much more complex than its validation). In addition, we harnessed GPT4 (OpenAI, 2023) to serve as the judge. Similar techniques to generate and validate solutions have been proposed in the context of code generation (Haluptzok et al., 2022). As the final output, BetaDescribe provides a set of possible descriptions ranked by their likelihood.

**Figure 1:**
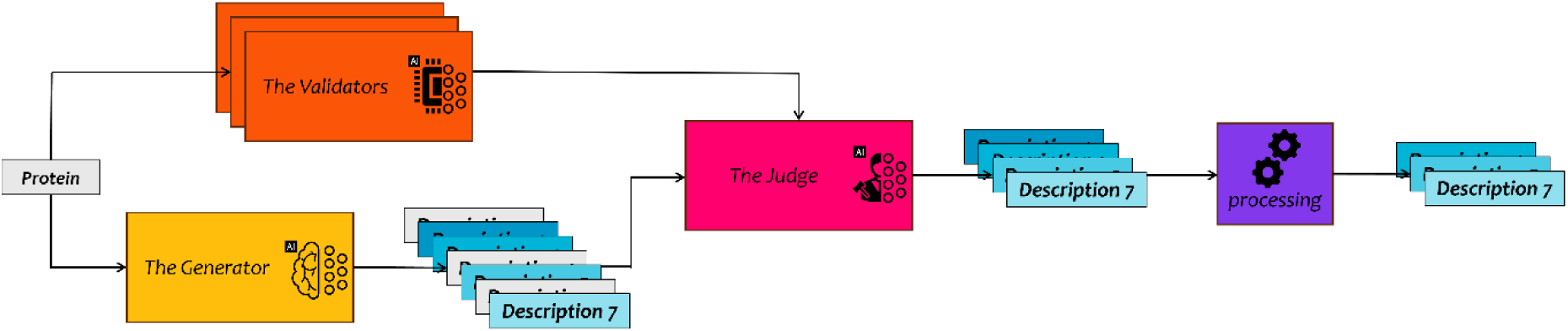
BetaDescribe workflow. The generator processes the protein sequences and creates multiple candidate descriptions. Independently, the validators provide simple textual properties of the protein. The judge receives the candidate descriptions (from the generator) and the predicted properties (from the validators) and rejects or accepts each description. Finally, BetaDescribe provides up to three alternative descriptions for each protein.

### The Generator

The generator is the central model in BetaDescribe. It is a decoder-only model (Radford et al., 2018) trained to generate textual descriptions of proteins in English, bridging the protein and English domains. The starting point of the model was LLAMA2, which was pretrained on two trillion English tokens by Meta (Touvron et al., 2023). Next, we continued training the model on 120 billion additional tokens, which include protein sequences and their descriptions, thus incorporating biological knowledge. The model was trained to predict the next token, i.e., given a sequence prefix, the model was trained to complete it. Training included several phases designed to gradually move from general English language modeling to generating textual descriptions of proteins.

#### Training

Starting from the original LLAMA2 model (Touvron et al., 2023), pretrained on general English text only (from multiple public open sources), we trained the generator in three consecutive stages. We first added protein sequences to the training data. These training data (Dataset 1, ∼2.9 × 10^10^ tokens) contained a mixture of English sentences from the RedPajama (https://www.together.ai/blog/redpajama) dataset (30%) and a large set of diverse proteins (70%), randomly sampled from the Uniref-90 dataset (Suzek et al., 2015). Training on both English and protein sequences together allowed the model to incorporate biological knowledge without losing the ability to generate English text.

While Dataset 1 contained both English and protein sequences, the English text was not directly connected to the protein sequences. In Dataset 2, we incorporated protein sequences and their cognate textual description (in English), thus adding expert vocabulary and knowledge regarding protein functionality. Specifically, this dataset contained protein sequences and their English descriptions (45%), English descriptions and their cognate proteins (45%), as well as English sentences not related to biology (10%). In total, this dataset contained ∼1.3 × 10^10^ tokens, in which the biological proteins and their cognate-rich textual description were derived from the UniProt dataset (The UniProt Consortium, 2017). For example, protein “A0A0V0V610” with the following protein sequence, “MGREDKTTWKSNYFLKLV[…]”, is trained to predict the following: “*FUNCTION$ Ribosomal protein P0 is the functional equivalent of E.coli protein L10, SIMILARITY$ Belongs to the universal ribosomal protein uL10 family[…]*”, and vice versa, i.e., the model is trained to predict the protein sequence given the rich description.

Finally, we trained the model to predict the protein description given the protein sequences (Dataset 3). This dataset contained ∼8.3 × 10^10^ tokens. Between the different stages, we used transfer learning, i.e., the optimal model trained in the previous stage was the starting point for the training of the next stage (Tan et al., 2018). We tested two pretraining models and different values for the learning rate for biological data processing. For the specific models, training hyperparameters, and dataset configurations, see Supplementary Information S1.

#### Multiple Candidate Descriptions

We used two techniques to generate multiple candidate descriptions. The first relies on changing the prompts used as input for the LLMs. For English LLMs, the prompt includes the instructions (the task, e.g., “shorten the following text”), and the context (additional information needed to perform the task, e.g., the text to be shortened). Different prompts lead to different responses (Sahoo et al., 2024). For our generator, we used three alternative prompts (Supplementary Information S2). Note that detailed instructions are not needed in our case, as the model was trained for the specific task of describing the function of the query protein.

For each prompt, we generated 15 alternative descriptions using a temperature hyperparameter (Ackley et al., 1985; Hinton et al., 2015). Generating text is done by choosing the next token until reaching the special token marking the end of the text or reaching a predefined maximum length. By default, the next token chosen is the one with the highest probability. However, the model can choose the next token based on the distribution of token probabilities. A high temperature flattens this distribution, thus introducing variability in the generated outputs, i.e., the descriptions (Supplementary Information S2).

#### Memorization and Generalization

Multiple proteins in the dataset have the same descriptions, which led the generator to memorize some of them. With a temperature of 0, most of the generated descriptions appear in the training set. This resembles a “search” operation within the description domain. The generation of novel descriptions could provide additional insights. Increasing the temperature hyperparameter allows the generator to predict descriptions that are not present in our training set. In a preliminary analysis, a temperature of 1.0 balanced description diversity and lack of hallucinations (not shown).

#### Architecture

The model architecture is based on the LLAMA2model with its default tokenization (Touvron et al., 2023). It is a decoder-only model with seven billion parameters, comprising 32 layers and 32 attention heads. The hidden state size is 4,096. See Touvron et al. (2023) for additional details on the architecture.

### The Validators

We trained three different validators, each predicting a specific protein property given the input protein sequence. We selected properties that can be accurately predicted and are relevant for many proteins. The three properties were: (1) higher-level taxonomic classification “viruses”, “bacteria”, “archaea” and “eukaryota” based on the UniProt lineage property; (2) subcellular localization consisting of 388 categories, including “acrosome”, “chlorosome envelope”, “Golgi apparatus lumen”, and “plasmodesma”. Since proteins may function in more than one location, this classification is multi-labeled; and (3) presence of enzymatic activity (a binary-classification task). The starting point for each validator is the ESM2 base model (Lin et al., 2023), which is associated with 150 million parameters. For training and evaluating the validators, we extracted the proteins and their corresponding labels from the UniProt dataset (The UniProt Consortium, 2017). For the models, tokenizers, training hyperparameters, and datasets, see Supplementary Information S3.

### The Judge

The judge determines the congruence between the validators’ predictions and a candidate’s description. For example, the judge is expected to reject a bacterial protein (property predicted by a validator) with a description that includes activity related to the eukaryotic spliceosome (generated by the generator). The judge is an external LLM trained on English text, some of which relate to biological knowledge. The generator and the validators convert the protein sequences to English descriptions, and English properties, respectively, and the resulting text is used as input for the judge (Figure 1). Specifically, we used a combination of rule-based decision and prompt-based queries to GPT4 (OpenAI, 2023) to reject unlikely descriptions (Supplementary Information S4).

### Selecting a Subset of Diverse Descriptions

From the resulting descriptions, we aim to select a subset of representative descriptions reflecting the diversity of the suggested protein functions. Specifically, we selected three descriptions from the descriptions that passed the judge (45 descriptions, 15 for each of the three prompts). To this end, we created a graph, in which nodes are descriptions, and edges are the string-based distances between two descriptions; specifically, we used the Character n-gram F score (ChrF; Popović, 2015; Supplementary Information S5). Next, we computed communities (clusters), i.e., groups of nodes (descriptions) that are more densely connected to each other than to the rest of the nodes (Blondel et al., 2008; Radicchi et al., 2004). We focused on the three largest communities, and for each such community, we selected the representative description with the highest average ChrF value. The yielded descriptions are ranked by the community size. We used the Networkx library to implement the graph and the search for communities (Hagberg et al., 2008).

### Evaluation

To evaluate the performance of BetaDescribe on the test set, we compared each inferred description to the true one, the latter provided by UniProt (The UniProt Consortium, 2017). Four alternative metrics were used to evaluate performance. First, the exact match binary metric assigns a value of 1 if the true and inferred descriptions are the exact same string, and 0 otherwise. Next, we utilized two metrics commonly used for evaluating machine-translated text: the ChrF score (Popović, 2015) and BLEU (Papineni et al., 2002), as implemented in SacreBLEU (Post, 2018). These metrics assign a score between 0 and 1, indicating the degree of similarity between the strings based on shared characters and words. Furthermore, we included the cosine similarity metric, which measures the distance between the embeddings of the descriptions, i.e., the numerical representations of the descriptions processed by the generator. Unlike the three previous string-based approaches, cosine similarity assesses the semantic meaning of the descriptions (Supplementary Information S5 provides detailed metric explanations).

## Results

### BetaDescribe Performance

A total of 2.5 million proteins comprising the test set were divided into three categories according to their similarity to the training set, evaluated by BlastP E-values (Altschul et al., 1990). Category 1 sequences lack BlastP hits (E-value > 10, 189 proteins), Category 2 sequences had statistically insignificant hits against the training data (1 < E-value <= 10, 172 proteins), and Category 3 sequences had nearly significant or significant hits (E-value <= 1, ∼ 2.5 × 10^6^ proteins). In our analysis, we compared BetaDescribe predictions to the best BlastP hit (i.e., the one with the lowest E-value) when searching the training set.

#### Providing Descriptions when BlastP is Unavailable

The performance of BetaDescribe on Category 1 proteins is provided in Table 1a. Within Category 1, the first prediction of BetaDescribe had an exact match to the true description for 27 out of the 189 proteins (14.3%). These exact descriptions were surprising because we demanded that no protein be shared between the training and the test datasets. To gain further insights into such cases, we inspected specific cases. An example of such a case is protein B0M8U4 in the test set, corresponding to the short peptide of “TDRNFLRL”. We found that a different protein in the training set, B0M3D0, shares the exact same sequence and description: “FMRFamides and FMRFamide-like peptides are neuropeptides”. BlastP failed in this case because BlastP searches for significant hits, and these peptides are too short to yield any E-values. Altogether, we found 38 cases with identical sequences (and identical or nearly identical descriptions) for a protein in the training set and another in the test set. Accordingly, we excluded all proteins with an identical sequence in the training and test data from all further analyses. The performance of Category 1 without those sequences is provided in Table 1b. In general, across all accuracy measures, the differences between the first, second, and third predictions were relatively small (Table 1).

**Table 1.**
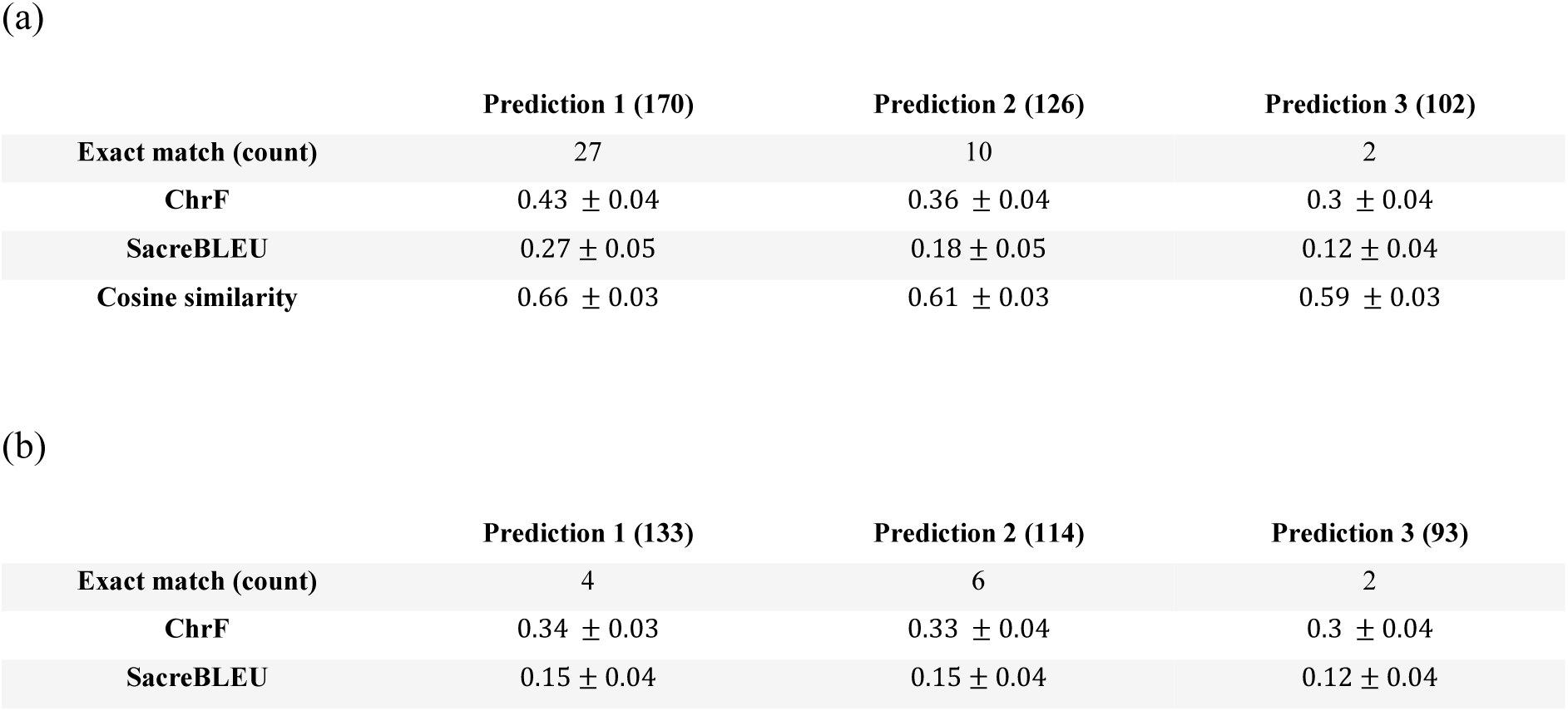
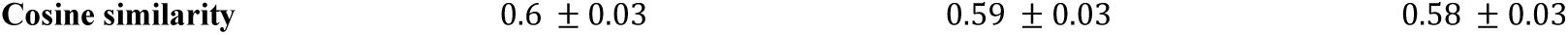
Performance of BetaDescribe on Category 1 proteins, i.e., test proteins without BlastP hits when searched against the training data. The number of descriptions for each column is stated in parentheses. (a) The performance considering the 189 proteins in Category 1. (b) The performance without the 38 proteins with identical sequences in the training data. Predictions 1,2, and 3 are the first, second, and third descriptions, respectively provided by BetaDescribe.

For the remaining 151 proteins, 69 proteins had the exact same description in the training set but for a protein with a different sequence. For example, protein A0A8C0XGC0 in the test set is 50 amino-acid long and has the following description in UniProt: “Protamines substitute for histones in the chromatin of sperm during the haploid phase of spermatogenesis. They compact sperm DNA into a highly condensed, stable and inactive complex”. Our model perfectly predicted this function, probably because the function exists in the training set. However, the same functional description appears for a related protein that differs from A0A8C0XGC0 by three one-character insertions or deletions (indels), and nine substitutions, i.e., 77% identity. BlastP failed to detect significant sequence similarity probably because of the presence of low-complexity regions and repetitive elements (Supplementary Information S6).

Notably, even when the performance scores are substantially less than 1, components of the predicted descriptions may be accurate. Consider the case of protein A0A1E3Q8Q4, for which the accuracy was 0.35, 0.081, and 0.61 for The ChrF, SacreBLEU, and cosine similarity scores, respectively. The description of this 84 amino-acid long protein in UniProt is:

“*FUNCTION$ Binds tightly to hydroxyapatite. Appears to form an integral part of the mineralized matrix. Probably important to cell-matrix interaction. Promotes Arg-Gly-Asp-dependent cell attachment, SUBCELLULAR LOCATION$ Secreted.*”

In comparison, our model prediction is:

“*FUNCTION$ Plays a role in cell adhesion and tissue remodeling. May be a cell-cell adhesion protein with cell-adhesive properties, SUBCELLULAR LOCATION$ Secreted, extracellular space, extracellular matrix, PTM$ May be proteolytically cleaved, SUBUNIT$ Interacts with SPRED1, SIMILARITY$ Belongs to the invertebrate Chitin-binding protein family.*”

The predicted description correctly captures that this protein is secreted and is involved in cell-cell and cell-extracellular matrix interactions. Here, too, BlastP’s failure to find a similar sequence in the training set is likely because of the presence of low-complexity regions (Supplementary Information S6), while a similar description appears in the training data. To conclude, for many of the proteins in this category, BlastP technically fails to find a hit as the proteins are short, include low-complexity regions, or both (Supplementary Information S7).

#### Providing Descriptions when BlastP E-value is High

Category 2 includes 172 proteins with no significant hits in the training data (E-value higher than 1). 21 of those proteins were very short and thus resembled some of the cases in Category 1, i.e., BlastP failed to find the exact matched protein in the training set because of the short length (Supplementary S7). For four of the remaining 151 proteins, BetaDescribe failed to provide a valid description (2.6%). Of the remaining 147 cases, BetaDescribe provided an exact match for two and three proteins for predictions 1, and 2, respectively. Although a Blast E-value above 1 is considered insignificant (usually the threshold is much lower, e.g., Moreno-Hagelsieb & Latimer, 2008), retrieving the description from the best hit (lowest E-value) returned five proteins with the exact match. When comparing the cosine similarity score, BetaDescribe performance (prediction 1) was superior to BlastP: 0.58 and 0.48, respectively (paired t-test; *p* < 0.0001; Supplementary Information S8). The ChrF and the SacreBLEU scores were not significantly different.

The following example highlights the benefits of providing several alternative descriptions. In the test set, the description of protein C6Y4A7 is:

“*FUNCTION$ The EMC seems to be required for efficient folding of proteins in the endoplasmic reticulum (ER), SUBCELLULAR LOCATION$ Endoplasmic reticulum membrane, SUBUNIT$ Component of the ER membrane protein complex (EMC), SIMILARITY$ Belongs to the membrane magnesium transporter (TC 1.A.67) family.*”

In this specific case, the third prediction of BetaDescribe was the most accurate (cosine similarity of 0.79): “*FUNCTION$ Part of the endoplasmic reticulum membrane protein complex (EMC) that enables the energy-independent insertion into endoplasmic reticulum membranes of newly synthesized membrane proteins. May be involved in Mg(2+) transport, SUBCELLULAR LOCATION$ Endosome membrane, SIMILARITY$ Belongs to the membrane magnesium transporter (TC 1.A.67) family.*”

The first and second predictions captured some aspects of the “true” protein description (ER-related function). The first (cosine score of 0.72) was:

“*FUNCTION$ Involved in the early part of the secretory pathway, SUBCELLULAR LOCATION$ Golgi apparatus membrane, SIMILARITY$ Belongs to the KISH family.*”, and the second (cosine score of 0.78):

“*FUNCTION$ Required for the assembly of the V0 complex of the vacuolar ATPase (V-ATPase) in the endoplasmic reticulum, SUBCELLULAR LOCATION$ Endoplasmic reticulum membrane, SIMILARITY$ Belongs to the VMA21 family.*”

The BlastP-based prediction (best hit had an E-value of 6.5) had a lower cosine score (0.45) and substantially deviated from the provided descriptions: “*FUNCTION$ Catalyzes the deamination of dCTP to dUTP, CATALYTIC ACTIVITY$ dCTP + H(+) + H2O = dUTP + NH4(+), PATHWAY$*

*Pyrimidine metabolism; dUMP biosynthesis; dUMP from dCTP (dUTP route): step 1/2, SUBUNIT$ Homotrimer,SIMILARITY$ Belongs to the dCTP deaminase family.*”

There is no clear value of a cosine score above which a prediction is considered reliable. From our experience, prediction scores above 0.6 were relatively accurate. With this cutoff, out of the 147 predictions, Prediction 1 of BetaDescribe was accurate in 55 predictions, and BlastP was accurate in 27 cases (Figure 2). However, in some of these 27 cases, the description yielded by BlastP was more accurate than all three predictions of BetaDescribe (Supplementary Information S9).

**Figure 2:**
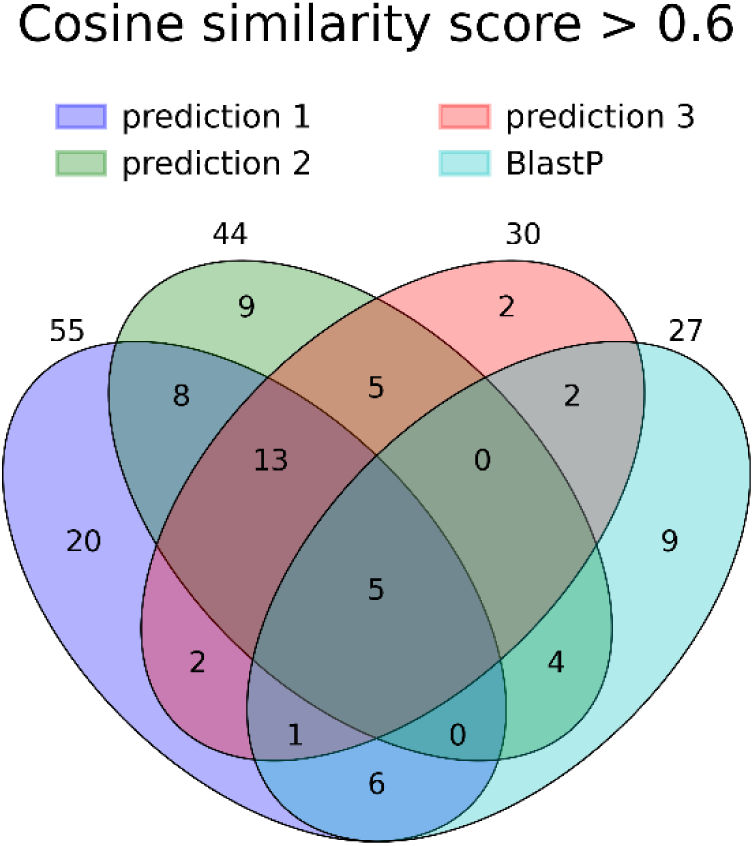
The Venn diagram displays the count and overlap of proteins with high cosine similarity scores (above 0.6). For example, five proteins achieved high cosine similarity scores for the three BetaDescribe predictions and the BlastP-based description. Additionally, 20, 9, 2, and 9 proteins had high cosine similarity scores exclusively in predictions 1, 2, 3, or using BlastP, respectively. After normalization, these counts correspond to 37.4%, 40.7%, 34.5%, and 18.3%, respectively.

#### Providing Descriptions when BlastP Hits are Significant

We sampled 1,000 proteins from the test set, for which similar proteins are present in the training set, i.e., E-values lower (or equal) to 1.0 when running BlastP against the training set. Taking the description from the closest hit as the predicted function yielded accurate predictions (Supplementary Information S8). Across all metrics, BlastP-based descriptions were significantly better than the ones provided by BetaDescribe (paired t-test; p < 0.0001), suggesting that functionality transfer based on BlastP between closely related proteins is highly accurate. However, we found a strong correlation between the accuracy of the BlastP prediction if it “agrees” with the BetaDescribe prediction. We divided the proteins by their E-value score into ten categories (one category of 235 proteins with a hit E-value of 0, and nine other bins with an equal number of proteins, 75 or 76). For each category, we divided the proteins into high and low values based on the cosine similarity score between the BlastP prediction and Prediction 1 of BetaDescribe. Figure 3 reports the average cosine similarity score of the BlastP hit. When the BlastP and BetaDescribe predictions are congruent, i.e., a cosine similarity score above the median for that bin, the BlastP predictions are more accurate compared to the case where BlastP and BetaDescribe predictions are incongruent (t-test; *p* < 0.001).

**Figure 3:**
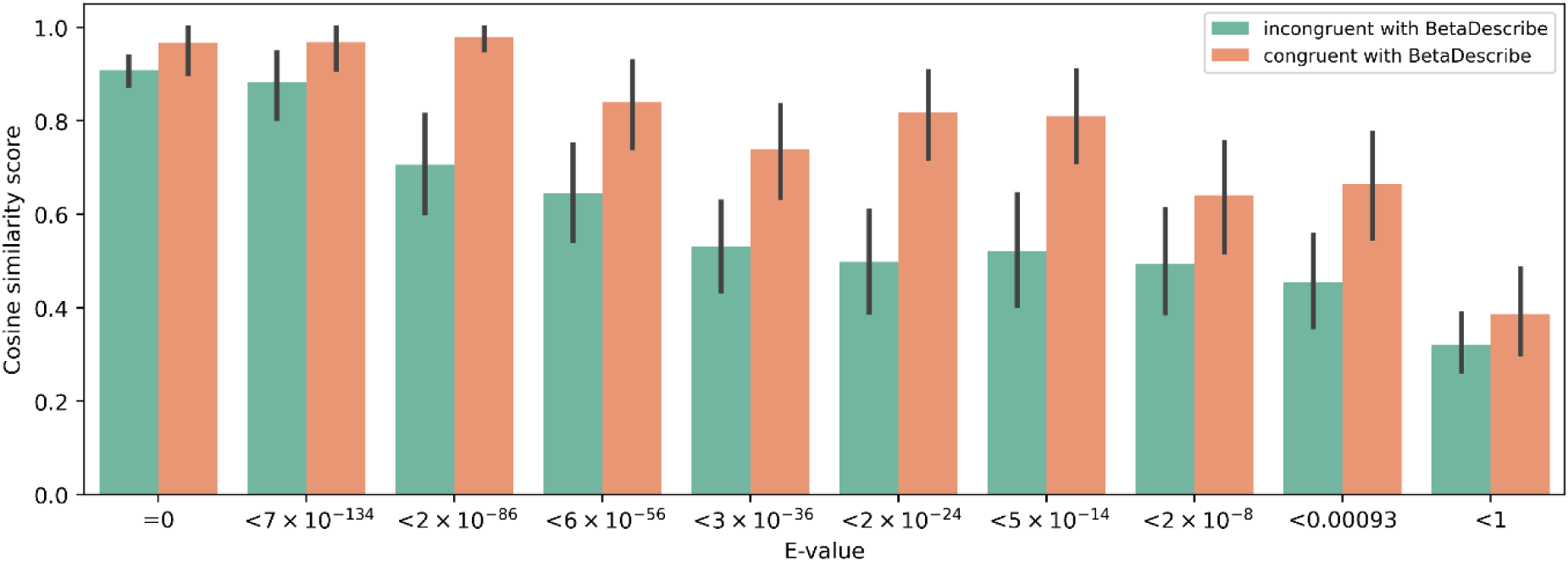
BlastP-based predictions are more accurate when congruent with BetaDescribe’s predictions. BlastP scores on Category 3 proteins, as a function of their E-value score as yielded by BlastP. In each bin, we divided the BlastP predictions into two groups: those that are congruent with BetaDescribe’s prediction 1 (cosine similarity score above the median) and those that do not.

#### Analyzing the Source of Errors

In some cases, the judge rejects a description generated by the generator based on input from the validators (Figure 1). Such cases can result from three scenarios: either the generator’s description is false, and the validators correctly disagree with it, or the generator’s description is correct, but the validator’s predictions are wrong, which is the source of the disagreement. Another option is that both the validator and the generated description are correct, but the judge erroneously determines they are incompatible. This last case occurs in 19% (see Supplementary Information S10 for detailed analyses).

### Public LLM Predictions for Protein Functions

Next, we evaluated the performance of predicting protein functions using public LLMs. We tested three LLMs trained on general knowledge: GPT4 (OpenAI, 2023), Gemini (Gemini Team et al., 2024), and Claude (https://www.anthropic.com/news/claude-3-5-sonnet). We asked these LLMs to predict the function of 30 test proteins from Category 3. In the prompt, we provided the protein sequence, as well as three examples for the required output (Supplementary Information S11). The cosine similarity score of the prediction and the true description was comparable among the three LLMs, with Claude, GPT4, and Gemini values of 0.49, 0.45, and 0.43, respectively. GPT4 and Gemini scores were not significantly different from scores obtained when the description for a given protein was randomly sampled from the training set (*p* = 0.37, 0.89, respectively). For comparison, BetaDescribe’s cosine similarity score was 0.86. For only two proteins, the public LLMs provided a cosine similarity score above 0.6: A0A0D2C110 and A0A0D6M674. The highest cosine similarity score was predicted by Claude for protein A0A0D2C110, with the “true” description:

“*FUNCTION$ Has a role in the initiation of DNA replication. Required at S-phase checkpoint, SUBCELLULAR LOCATION$ Nucleus, SIMILARITY$ Belongs to the SLD2 family.*”

For this protein Claude provided the following description: “*FUNCTION$ Transcriptional regulator involved in chromatin remodeling and gene expression regulation, particularly during development and cellular differentiation, SUBCELLULAR LOCATION$ Nucleus, SIMILARITY$ Belongs to the ARID (AT-rich interaction domain) family of DNA-binding proteins*”

### Providing Descriptions for Proteins with Unknown Functions

We demonstrate the usage of BetaDescribe by analyzing six examples of proteins with no experimentally proven functionality but whose function can be predicted by other attributes, e.g., location in the genome and structural features. Specifically, we selected five proteins from three different viruses, encompassing two predicted envelope glycoproteins and three RNA-depended RNA polymerase subunits. We also analyzed a newly discovered bacterial protein involved in the immune system.

#### TGV-S

The first example is the putative spike (S) protein of the recently identified nidovirus, the Trout Granulomatous Virus (TGV). The inferred function of this protein is based on the genomic localization of its open reading frame (ORF) and the protein’s domain structure, which are typical for nidoviruses (Karniely et al., 2023). BlastP search with TGV-S sequence against the training set yielded a significant hit (Q28042; E-value < 10^−5^). This known protein has an unrelated function in fertilization or embryonic development. In contrast, BetaDescribe provided two valid predictions, each describing a viral envelope protein (Table 2).

**Table 2:**
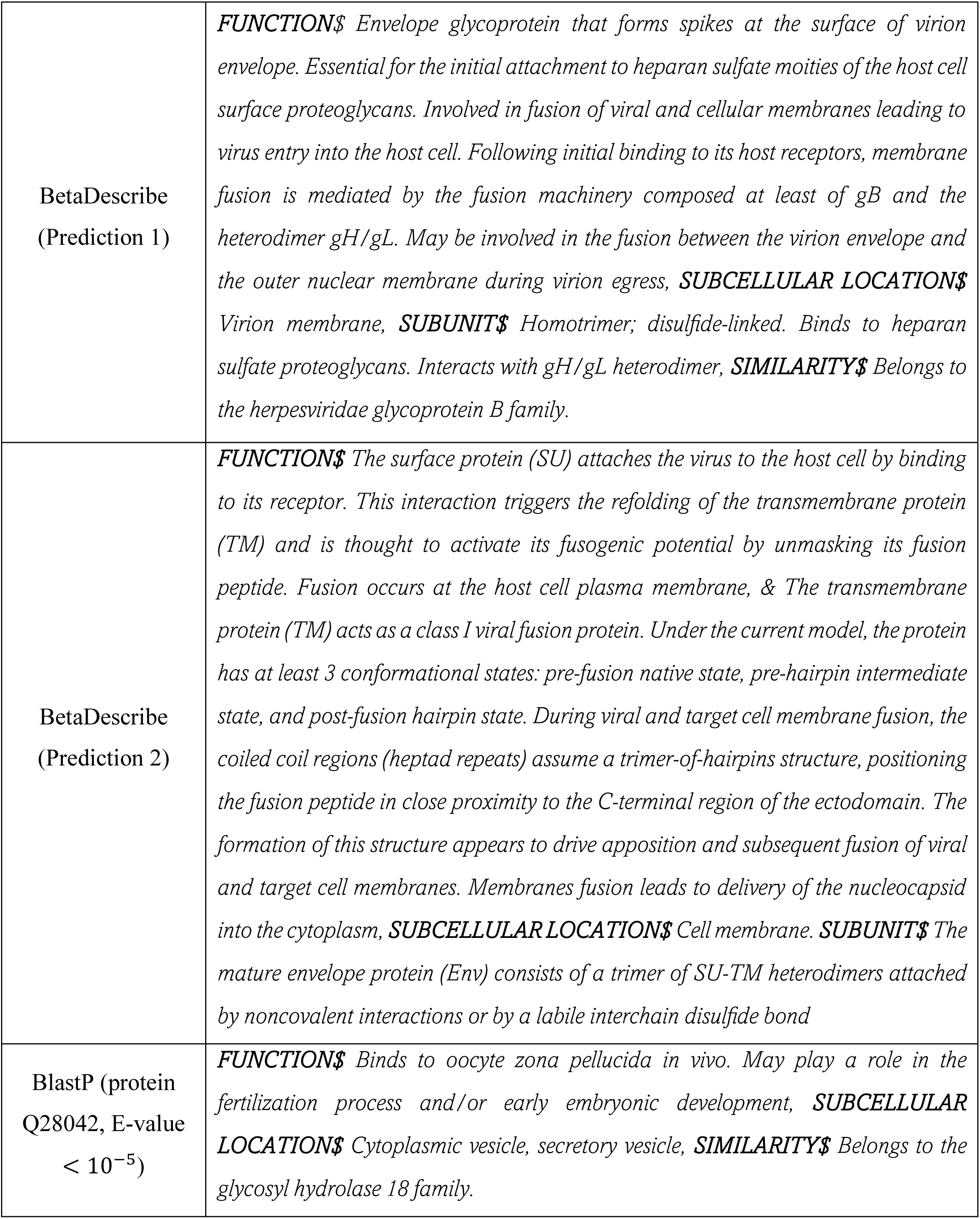
Prediction for the TGV-S protein by BetaDescribe (Predictions 1 and 2) and BlastP.

#### SnRV-Env

The second example is the envelope (Env) protein of the snakehead retrovirus (SnRV). The genome of this fish virus was sequenced about 30 years ago (Hart et al., 1996), but no experimental data support the functionality of this protein. The SnRV-Env functionally is predicted by the genomic localization of the *env* gene, typical to retroviruses (downstream to the *gag-pol* ORFs), and an inferred leader peptide and a transmembrane domain (Hart et al., 1996). Most retroviral Env proteins are targeted to the plasma membrane and are essential for receptor binding, membrane fusion, and viral entry into the host cells. Querying the protein sequence against the training set yielded a hit with an E-value of 0.5 to a protein, A0A0R1ZYJ7, with an unrelated function of pseudouridine synthesis. BetaDescribe provided two predictions: the first inferred functions related to viral envelope proteins, and the second predicted membranal localization (Supplementary Information S12).

#### Descriptions for Three TiLV Proteins

Tilapia lake virus (TiLV) is a negative-stranded RNA virus first identified in 2014 in northern Israel (Eyngor et al., 2014). The virus infects both wild and farmed tilapia populations. Since its discovery, TiLV has been detected in various regions across Asia, Africa, and South America, threatening the food security of millions of people (Jansen et al., 2019). By and large, when discovered, TiLV’s ten main proteins showed no significant sequence similarity to other known viral proteins (Bacharach et al., 2016). Among these ten, Proteins 1 – 3 were predicted to serve as subunits of TiLV RNA-depended RNA polymerase (RdRP) (Abu Rass et al., 2022), a prediction recently validated experimentally and structurally (Arragain et al., 2023). These sequences and their associated functions were not included in the training or the test data. Among these three proteins, the predictions based on BlastP correctly identified only Protein 1 as polymerase (Supplementary Information S12). In contrast, BetaDescribe correctly assigned polymerase activity for all three proteins. For Protein 1, the top prediction of BetaDescribe correctly assigned RdRP activity, while for Proteins 2 and 3, BetaDescribe top predictions accurately assigned a polymerase activity, but inaccurately described the activity to be DNA-dependent. Of note, some of BetaDescribe’s alternative predictions are likely to be false, e.g., the proteasome connection in the second prediction for Protein 1 (Supplementary Information S12). Interestingly, an endonuclease-like domain was identified in Protein 3, but its functionality remains elusive (Arragain et al., 2023); such activity is included in an alternative BetaDescribe prediction.

#### H5TRP0

To further demonstrate the utility of BetaDescribe’s ability in analyzing non-viral proteins, we provided a query for the bacterial protein H5TRP0. We selected this protein because its function was described only recently after the training and test datasets were created. This protein, termed Cas12m, operates within the Clustered Regularly Interspaced Short Palindromic Repeats (CRISPR) system (Bigelyte et al., 2024), which is an adaptive immune system in prokaryotes that targets foreign DNA or RNA sequences with high specificity (Ishino et al., 1987). Searching for the most related protein in the training set yielded an insignificant hit to an unrelated protein with serine protease activity (A0A916UE10, E-value of 1). In contrast, BetaDescribe generated two out of three predictions that correctly linked it to CRISPR activity (Table 3). Although the third description is not directly Cas-associated (but transposition-related), recent studies reveal that the RNA-guided DNA binding and cleavage activity of Cas12 originates from transposon-encoded nuclease TnpB. This nuclease promotes transposon survival and spread and performs similar reactions to Cas12 (Wiegand et al., 2024).

**Table 3:** Prediction for the TGV-S protein by BetaDescribe (Predictions 1 - 3) and BlastP.

|  |  |
| --- | --- |
| BetaDescribe<br>(Prediction 1) | **FUNCTION\$** *CRISPR (clustered regularly interspaced short palindromic repeat) is an adaptive immune system that provides protection against mobile genetic elements (viruses, transposable elements and conjugative plasmids). CRISPR clusters contain spacers, sequences complementary to antecedent mobile elements, and target invading nucleic acids. CRISPR clusters are transcribed and processed into CRISPR RNA (crRNA).* |
| BetaDescribe<br>(Prediction 2) | **FUNCTION\$** *CRISPR (clustered regularly interspaced short palindromic repeat) is an adaptive immune system that provides protection against mobile genetic elements (viruses, transposable elements and conjugative plasmids). CRISPR clusters contain sequences complementary to antecedent mobile elements and target invading nucleic acids. CRISPR clusters are transcribed and processed into CRISPR RNA (crRNA),*<br>**COFACTOR\$** *Mg(2+) or Mn(2+) required for ssDNA cleavage activity., & iron-sulfur cluster,*<br>**SIMILARITY\$** *Belongs to the CRISPR-associated exonuclease Cas4 family.* |
| BetaDescribe<br>(Prediction 3) | **FUNCTION\$** *Involved in the transposition of the insertion sequence IS5,*<br>**SIMILARITY\$** *Belongs to the transposase 11 family.* |
| BlastP (protein<br>A0A916UE10, E-value of 1) | **FUNCTION\$** *ATP-dependent serine protease that mediates the selective degradation of mutant and abnormal proteins as well as certain short-lived regulatory proteins. Required for cellular homeostasis and for survival from DNA damage and developmental changes induced by stress. Degrades polypeptides processively to yield small peptide fragments that are 5 to 10 amino acids long. Binds to DNA in a double-stranded, site-specific manner,*<br>**CATALYTIC ACTIVITY\$** *Hydrolysis of proteins in presence of ATP.,*<br>**SUBUNIT\$** *Homohexamer. Organized in a ring with a central cavity,*<br>**SUBCELLULAR LOCATION\$** *Cytoplasm,*<br>**INDUCTION\$** *By heat shock,*<br>**SIMILARITY\$** *Belongs to the peptidase S16 family.* |

### Identifying Functionally Important Protein Regions

Functionally important protein regions can be identified through mutagenesis experiments (Hutchison et al., 1978). Next, we tested whether BetaDescribe could be used to identify functionally important regions. To this end, we performed *in-silico* alanine-scanning mutagenesis experiments and tested the effect of these alterations on the description. Specifically, we quantified the fit (negative log-likelihood) of the description of the wild-type protein to that of the altered protein sequence, expecting that disturbing functionally important regions would substitutionally reduce this fit (Supplementary Information S13).

We report preliminary results for this approach on the extensively studied human insulin protein (P01308; an additional example, the protein RecA, is provided in Supplementary Information S13). The preproinsulin precursor is comprised of four domains: Signal peptide, Insulin B chain, C peptide, and Insulin A chain. The Signal peptide and the C peptide (which are less evolutionary conserved) are excised during insulin protein maturation, leaving insulin B and A chains (which are highly evolutionary conserved) to form functional insulin (Steiner et al., 2009). Figure 4 reports the importance of each amino acid in the insulin sequence, deduced from the change in the fit of the description of the wild-type protein to that of the mutated protein sequence. As expected, BetaDescribe descriptions were mostly affected by mutating amino acids within the Insulin A and B chains, and substantially less by mutations in the Signal peptide and C peptide. This analysis suggests that the BetaDescribe model learned to capture biological meaningful domains, which can be used to infer them.

**Figure 4:**
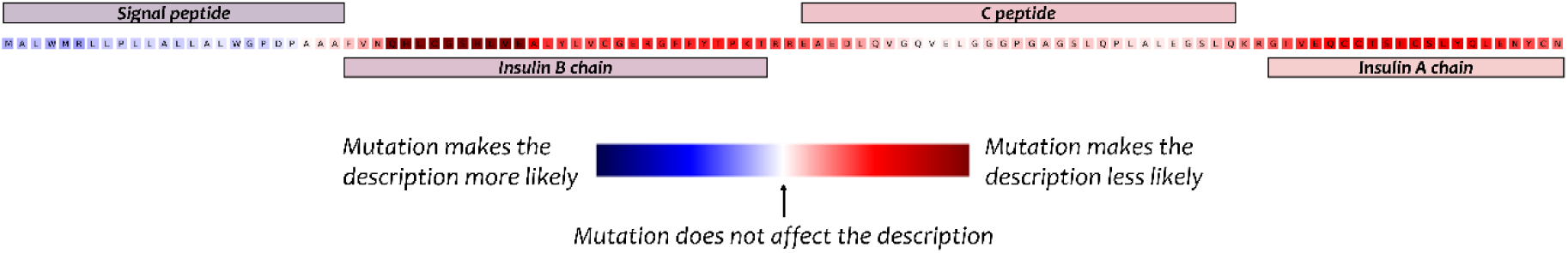
Identifying functionally important regions for the preproinsulin protein. The four main regions of the insulin are marked: Signal peptide, Insulin B chain, C peptide, and Insulin A chain.

## Discussion

BetaDescribe harnesses the generative capabilities of LLMs to provide rich and accurate textual descriptions for any protein of interest. To this end, the generator was trained on rich datasets of millions of biological pairs of sequences and their descriptions. In this work, we provide evidence that BetaDescribe is mostly useful in cases where BlastP fails (Category 1) or yields insignificant hits (Category 2). BetaDescribe provides up to three alternative possible predictions for each protein. In addition, as exemplified by Category 3 proteins, when the predictions of BetaDescribe and BlastP are congruent, the confidence in each of the provided predictions increases. Notably, disagreements among BetaDescribe alternative predictions suggest that each prediction is uncertain. Alternative descriptions could be viewed as hypotheses that need to be experimentally tested.

The relationship between BlastP and BetaDescribe resembles the one of traditional information retrieval (such as Google Search) and generative information retrieval (such as DSI transformer; Tay et al., 2022), which works by feeding a query into a neural network, and the network returns the document that matches this query. Accordingly, BetaDescribe allows an immediate and intuitive way to obtain descriptions for a given protein rather than a list of homologous proteins. Thus, BetaDescribe serves as a complementary tool for function prediction based on remote homology algorithms. In the future, we expect the users to be able to repeatedly communicate with such LLM-based predictors and influence the prediction by providing additional information, e.g., from unpublished experiments.

Explaining model predictions is common in the NLP domain (Atanasova et al., 2020). As exemplified by the preproinsulin and the RecA analysis (Supplementary Information S13), explanation techniques can be utilized to better understand the functional importance of different protein regions. In our analysis, we applied a simple sliding window alanine scanning approach for this task. Future work could investigate exposing interacting regions via *in-silico* complex mutagenesis. We envision a system that not only predicts protein functions but also highlights functionally important regions, providing context-specific insights into their roles.

Emerging deep-learning-based tools such as GPSFun (Yuan et al., 2024), ProteInfer (Sanderson et al., 2023), StarFunc (Zhang et al., 2024), and ESM3 (Hayes et al., 2024) classify proteins (or subdomains) to a predefined set of known functions, e.g., Gene Ontology (GO) or Enzyme Commission Identifier (ECI). While these tools predict from a selected predefined set of options, BetaDescribe aims to provide comprehensive English descriptions dedicated to explaining the function of the protein in question. BetaDescribe is an LLM trained in English and protein data. We speculate that LLM-based approaches will become central in computational tasks including protein engineering, evolutionary inference, and drug design. Looking ahead, we envision that the opposite direction, i.e., designing novel proteins based on rich descriptions of their intended function, will revolutionize and expedite protein-based research.

## Data Availability

Code and trained models are available at https://github.com/technion-cs-nlp/BetaDescribe-code.

## Acknowledgments

T.P. and Y.B. have received funding from the Israel Science Foundation (Grants 2818/21 and 448/20, respectively).

