## Supplementary Information for "Protein2Text: Providing Rich Descriptions for Protein Sequences"

### Supplementary Information S1: Generator – Training Hyperparameters and Training Datasets

#### *Preliminary Stage*

As a starting point for training the generator, we used the LLAMA2 model developed by Meta (Touvron et al., 2023). This model was trained on 2 trillion English tokens. The model was downloaded from Huggingface (Wolf et al., 2020). The training dataset is English only and is from multiple public open sources.

#### *Stage 1*

We downloaded the Uniref-90, which contains about 175 million protein sequences (Suzek et al., 2015), and the RedPajama dataset, which includes general English text (https://www.together.ai/blog/redpajama). From these datasets, we established the training data which includes 70% protein sequences (~ 4,900,000 proteins) and 30% English text (~ 2,100,000 sentences). The model architecture is the same as the LLAMA2 model, with the initial weights equal to those provided by the LLAMA2 model. We did some experiments to find the best learning rate and initial models (see below) and chose a learning rate of $3 \times 10^{-5}$ with the LLAMA2 model (Touvron et al., 2023).

Regarding the specific hyperparameters, we kept the same batch size (1,024) and the maximum sequence length (4,096) as was done in the training of the LLAMA2 model. Due to memory limitations, we applied the following techniques to reduce the memory footprint: numbers are kept in a Brain Floating Point (bfloat16) format (Kalamkar et al., 2019), gradient accumulation of 128 (with a micro-batch size of 1), and flash-attention 2 (Dao et al., 2022). We used a constant scheduler, i.e., a fixed learning rate for the entire training, and the AdamW optimizer (Loshchilov & Hutter, 2018), with betas of 0.9 and 0.999. The training was done with the Accelerator library (Wolf et al., 2020), on a single node with eight cores of NVIDIA A100-SXM4-80GB.

#### *Choosing Architecture and Learning Rate for Stage 1*

For optimal results, we started by finding the best model and learning rates for the biological data. We did experiments to evaluate the optimal learning rates as well as the initial model. We considered two models of similar sizes (about 7 billion parameters): Mistral (Jiang et al., 2023) and LLAMA2 (Touvron et al., 2023). Each of these models was trained for 500 training steps with four learning rates: $3 \times 10^{-5}$, $1 \times 10^{-5}$, $1 \times 10^{-6}$, and $1 \times 10^{-7}$. The models were evaluated on three different validation datasets: English (RedPajama), protein sequences (Uniref-90), and protein sequences and their descriptions (from UniProt). Table S1 reports the validation loss of the models trained with different learning rates. As expected, the loss of the biological data, and the protein sequences with their corresponding descriptions, starts high and decreases. However, the loss on the English dataset starts low and slightly increases. We anticipated this behavior as the model was trained for many steps on a similar English dataset. The performance of the LLAMA2 with a learning rate of $3 \times 10^{-5}$ had the lowest loss on the biological data, and thus, we decided to continue training with this specific learning rate.

*Table S1: We report the validation perplexity of LLAMA2 (a) and Mistral (b). For each model, we tested four different learning rates for 500 steps (corresponding to 2.1 billion tokens). Validation was conducted on three datasets: (1) English sentences (RedPajama); (2) protein sequences (Uniref-90); and (3) proteins and their corresponding descriptions (from UniProt). Training data contain 70% proteins and 30% English (Dataset 1).*

(a)

| LLAMA2 | | | | | | | |
|---|---|---|---|---|---|---|---|
| **Learning rate** | **Validation set** | **0** | **100** | **200** | **300** | **400** | **500** |
| $3 \times 10^{-5}$ | English | 1.7686 | 1.7788 | 1.7831 | 1.786 | 1.7883 | 1.7895 |
| | Proteins | 4.6591 | 4.3469 | 4.3245 | 4.3101 | 4.2963 | **4.2835** |
| | Descriptions | 2.827 | 2.8223 | 2.8074 | 2.8214 | 2.8005 | 2.7961 |
| $1 \times 10^{-5}$ | English | 1.7686 | 1.7618 | 1.7608 | 1.7605 | 1.7598 | 1.7597 |
| | Proteins | 4.6591 | 4.3465 | 4.3306 | 4.3216 | 4.3149 | 4.3026 |
| | Descriptions | 2.827 | 2.7885 | 2.789 | 2.7886 | 2.7834 | 2.7805 |
| $1 \times 10^{-6}$ | English | 1.7686 | 1.7665 | 1.7621 | 1.7612 | 1.76 | **1.7588** |
| | Proteins | 4.6591 | 4.3805 | 4.3643 | 4.3529 | 4.3451 | 4.3409 |
| | Descriptions | 2.827 | 2.7858 | 2.7799 | 2.7783 | 2.7762 | **2.7743** |
| $1 \times 10^{-7}$ | English | 1.7686 | 1.7682 | 1.7682 | 1.7709 | 1.7706 | 1.7688 |
| | Proteins | 4.6591 | 4.5394 | 4.4688 | 4.4228 | 4.4017 | 4.3937 |
| | Descriptions | 2.827 | 2.8155 | 2.8074 | 2.8051 | 2.7975 | 2.7936 |

(b)

| Mistral | | | | | | | |
|---|---|---|---|---|---|---|---|
| **Learning rate** | **Validation set** | **0** | **100** | **200** | **300** | **400** | **500** |
| $3 \times 10^{-5}$ | English | 1.8559 | 2.0395 | 2.1036 | 2.1301 | 2.1304 | 2.1513 |
| | Proteins | 4.8126 | 4.5226 | 4.4713 | 4.4426 | 4.418 | 4.3975 |
| | Descriptions | 2.8405 | 3.0433 | 3.0032 | 3.0125 | 2.9999 | 2.9918 |
| $1 \times 10^{-5}$ | English | 1.8559 | 1.8659 | 1.878 | 1.8895 | 1.8936 | 1.8983 |
| | Proteins | 4.8126 | 4.4834 | 4.4507 | 4.4299 | 4.4115 | **4.3928** |
| | Descriptions | 2.8405 | 2.848 | 2.8296 | 2.8273 | 2.828 | 2.814 |
| $1 \times 10^{-6}$ | English | 1.8559 | 1.821 | 1.8189 | 1.817 | 1.8153 | **1.8143** |

| | | | | | | | |
|---|---|---|---|---|---|---|---|
| | Proteins | 4.8126 | 4.4982 | 4.4744 | 4.4671 | 4.4573 | 4.4448 |
| | Descriptions | 2.8405 | 2.7822 | 2.7812 | 2.7833 | 2.7793 | **2.7769** |
| $1 \times 10^{-7}$ | English | 1.8559 | 1.8354 | 1.8298 | 1.8268 | 1.8247 | 1.8232 |
| | Proteins | 4.8126 | 4.5595 | 4.5403 | 4.5277 | 4.5167 | 4.5072 |
| | Descriptions | 2.8405 | 2.7994 | 2.7902 | 2.7863 | 2.7839 | 2.7829 |

#### *Stage 2*

We trained the model on a mixture of the UniProt (The UniProt Consortium, 2017) and the RedPajama datasets. Unlike the training in Stage 1, which contained sequences and English text, here the UniProt dataset contains both protein sequences and their English descriptions (32,409,736 pairs of proteins and English descriptions). We converted the UniProt data from a Json format to a text format, extracting only specific fields, specifically: function, catalytic activity, pathway, subcellular localization, domains, cofactors, PTMs, subunits, assignment to protein families, activity regulations, keywords, and features. An example of the resulting text for protein entry A0A6I7XUQ0 is:

protein sequence:
MTRIILPGKTIGIIGGGQLGRMMALAAKEMGYKIAVLDPTKHSPCAQVADIEIVASYDDLKAIQHLAEIS
DVVTYEFENIDYRCLQWLEKHAYLPQGSQLLSKTQNRFTEKNAIENAGLPVATYRLVQTQEQLTEAITE
LSYPSVLKTTTGGYDGKGQVVLRSEADVDKARKLANAAECILEKWVPFEKEVSVIVIRSVSGETKVFP
VAENIHVNNILHESIVPARITEELSQKAIAYARVLADELELVGTLAVEMFATADGEIYINELAPRPHNSG
HYTQDACETSQFGQHIRAICNLPLGETNLLKPVVMVNILGEHIEGVLRQVNRLTGCYLHLYGKEEAKA
QRKMGHVNILNDNIEVALEKAKSLHIWDHQEQLLEGKR description: FUNCTION$ Catalyzes the ATP-dependent conversion of 5-aminoimidazole ribonucleotide (AIR) and HCO(3)(-) to N5-carboxyaminoimidazole ribonucleotide (N5-CAIR), FUNCTION$ Catalyzes the ATP-dependent conversion of 5-aminoimidazole ribonucleotide (AIR) and HCO(3)- to N5-carboxyaminoimidazole ribonucleotide (N5-CAIR), CATALYTIC ACTIVITY$ 5-amino-1-(5-phospho-beta-D-ribosyl)imidazole + ATP + hydrogencarbonate = 5-carboxyamino-1-(5-phospho-D-ribosyl)imidazole + ADP + 2 H(+) + phosphate, PATHWAY$ Purine metabolism; IMP biosynthesis via de novo pathway; 5-amino-1-(5-phospho-D-ribosyl)imidazole-4-carboxylate from 5-amino-1-(5-phospho-D-ribosyl)imidazole (N5-CAIR route): step 1/2, SUBUNIT$ Homodimer, SIMILARITY$ Belongs to the PurK/PurT family. keywords: ATP-binding & Ligand, Ligase & Molecular function, Nucleotide-binding & Ligand, Purine biosynthesis & Biological process. features: Domain* 1, Binding site* 7.

We removed all proteins that do not have a specific functional property in their description. The training data at this stage were divided in a way that 45% (corresponding to 2,100,000 samples) of the data were assigned to generate the protein amino-acid sequence given the description, 45% of the data were assigned to generate the description given the protein sequence, and 10% (corresponding to 460,000 sentences) were assigned for English sentences. The same hyperparameters were used as described in Stage 1.

#### *Stage 3*

We trained the model to generate the descriptions given the amino-acid sequence from the UniProt dataset (The UniProt Consortium, 2017), containing 20,480,000 samples of proteins and their descriptions. The optimal model from Stage 2 was the starting position for this stage. We used the same training hyperparameters and only reduced the learning rate to $1 \times 10^{-5}$.

### Supplementary Information S2: Effect of Temperature and Multiple Prompts on Diversity

#### *The Temperature Hyperparameter*

Decoder-only models are trained to predict the next token, i.e., given the start of a sentence, the models are trained to predict the next word. Typically, the next predicted token is the one with the highest probability. However, it is possible to sample the next token from a distribution based on token probabilities. Low temperature results in a sharper distribution of tokens, i.e., the next token is likely to be the one with the highest probability. In contrast, high temperature flattens the distribution, increases diversity, and thus produces different alternative descriptions. We used the value of 1.0 for the temperature.

Formally, consider $t_i$, $l_i$, $P(.)$ to be the $i$'th token from a vocabulary with size $n$, the logits of the $t_i$ and the probability function, respectively. With the softmax approach, we select the token with the highest probability:

$$P(t_i) = \frac{e^{l_i}}{\sum_{k=1}^{n} e^{l_k}}$$

This expression is modified to account for the temperature, $t$, but here, we sample the next token according to the resulting probability:

$$P(t_i) = \frac{e^{l_i/t}}{\sum_{k=1}^{n} e^{l_k/t}}$$

#### *Alternative Prompts*

We considered three different prompts (Table S2): (1) protein sequence alone; (2) protein sequence with the "*FUNCTION$*" property; and (3) protein sequence with double space and the "*FUNCTION$*" property. For the above example, the three prompts are:

*Table S2: Detailed prompts used for generating the descriptions.*

| | |
|---|---|
| **Prompt 1** | protein sequence: MTRIILPGKTIGIIGGGQLGRMMALAAKEMGYKIAVLDPTKHSPCAQVADIEIVASYDDL KAIQHLAEISDVVTYEFENIDYRCLQWLEKHAYLPQGSQLLSKTQNRFTEKNAIENAGLP VATYRLVQTQEQLTEAITELSYPSVLKTTTGGYDGKGQVVLRSEADVDKARKLANAAE CILEKWVPFEKEVSVIVIRSVSGETKVFPVAENIHVNNILHESIVPARITEELSQKAIAYAR VLADELELVGTLAVEMFATADGEIYINELAPRPHNSGHYTQDACETSQFGQHIRAICNLP LGETNLLKPVVMVNILGEHIEGVLRQVNRLTGCYLHLYGKEEAKAQRKMGHVNILNDN IEVALEKAKSLHIWDHQEQLLEGKR description: |

| | |
|---|---|
| **Prompt 2** | protein sequence:<br>MTRIILPGKTIGIIGGGQLGRMMALAAKEMGYKIAVLDPTKHSPCAQVADIEIVASYDDL<br>KAIQHLAEISDVVTYEFENIDYRCLQWLEKHAYLPQGSQLLSKTQNRFTEKNAIENAGLP<br>VATYRLVQTQEQLTEAITELSYPSVLKTTTGGYDGKGQVVLRSEADVDKARKLANAAE<br>CILEKWVPFEKEVSVIVIRSVSGETKVFPVAENIHVNNILHESIVPARITEELSQKAIAYAR<br>VLADELELVGTLAVEMFATADGEIYINELAPRPHNSGHYTQDACETSQFGQHIRAICNLP<br>LGETNLLKPVVMVNILGEHIEGVLRQVNRLTGCYLHLYGKEEAKAQRKMGHVNILNDN<br>IEVALEKAKSLHIWDHQEQLLEGKR description: FUNCTION$ |
| **Prompt 3** | protein sequence:<br>MTRIILPGKTIGIIGGGQLGRMMALAAKEMGYKIAVLDPTKHSPCAQVADIEIVASYDDL<br>KAIQHLAEISDVVTYEFENIDYRCLQWLEKHAYLPQGSQLLSKTQNRFTEKNAIENAGLP<br>VATYRLVQTQEQLTEAITELSYPSVLKTTTGGYDGKGQVVLRSEADVDKARKLANAAE<br>CILEKWVPFEKEVSVIVIRSVSGETKVFPVAENIHVNNILHESIVPARITEELSQKAIAYAR<br>VLADELELVGTLAVEMFATADGEIYINELAPRPHNSGHYTQDACETSQFGQHIRAICNLP<br>LGETNLLKPVVMVNILGEHIEGVLRQVNRLTGCYLHLYGKEEAKAQRKMGHVNILNDN<br>IEVALEKAKSLHIWDHQEQLLEGKR description: FUNCTION$ |

If given Prompt 1, the first predicted token was not "*FUNCTION$*", we considered this prediction to be invalid. Of note, prompts 2 and 3 are always valid. For the first prompt, we achieved a validity score of 52.3%, 72.4%, and 81.2% for Categories 1, 2, and 3 (as defined in the main text), respectively.

#### *Combining the Temperature and the Alternative Prompts Provides Diversity in the Generated Descriptions*

The goal of accounting for the temperature and alternative prompts is to introduce stochasticity in the model, thus providing several descriptions for each input protein sequence. Specifically, we selected a temperature $t = 1.0$ and generated 15 alternative descriptions for each alternative prompt (totaling 45 alternative descriptions). After removing invalid predictions and those rejected by the judge, we plotted the shared and unique descriptions of each prompt. As can be seen in Figure S1, most alternative descriptions are unique, i.e., it is rare that the different prompts have the same exact prediction.

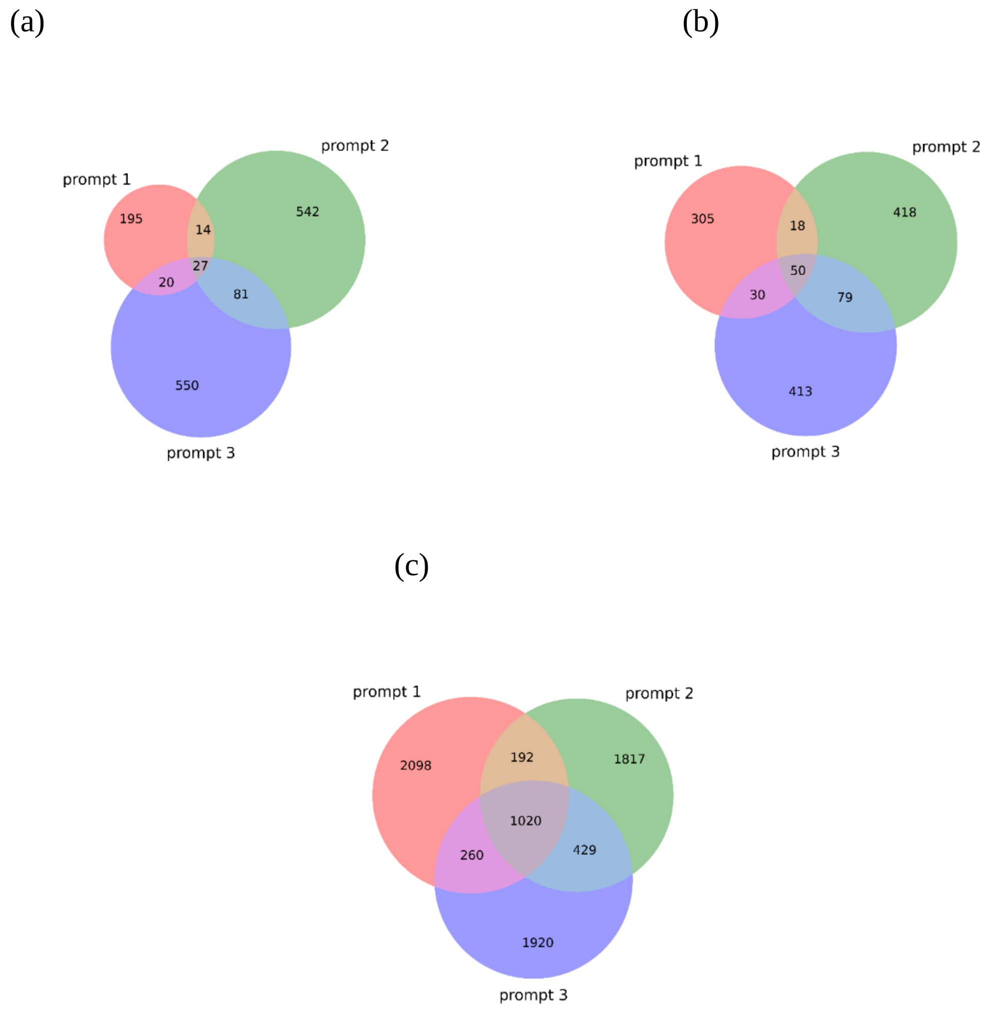


*Figure S1: We report the number of unique and shared descriptions using the different prompts on Categories 1, 2, and 3 for figures (a), (b), and (c), respectively. For example, in Category 1, we analyzed 151 proteins and obtained 6,795 descriptions (151 proteins times 45 alternative descriptions). After processing (verifying the function property), removing cases rejected by the judge, and removing duplications, we received the following 1,429 descriptions. For example, of these 1,429 descriptions, 14 generated by prompts 1 and 2 were identical but different from that generated in prompt 3.*

### Supplementary Information S3: Validators Architecture, Training, Datasets and Tokenization

We fine-tuned the ESM2 (Lin et al., 2023), an encoder-only model with 150 million parameters for predicting each of the following properties: subcellular localization, higher taxonomy, and enzyme activity. The base model has 30 hidden layers, 20 attention heads, a hidden size of 640, an intermediate size of 2,560, and a max sequence length of 1,026. The tokenizer for the model encodes each of the amino acids separately, and thus, the vocabulary size is 33 (including special tokens).

We used the following training parameters for each of the validators: maximum sequence length of 1,026 (same as the pretraining), batch size of 8, learning rate of $2 \times 10^{-5}$, 2,000 steps for warmup, weight decay of 0.01, and maximum training steps of 1,200,000. We trained with a mixed-precision approach (Micikevicius et al., 2018), i.e., parts of the model are saved in a reduced format using only 16 bits compared to the traditional training that uses 32 bits. We used the Huggingface library for this training (Wolf et al., 2020), and a single core of NVIDIA RTX A6000 48GB.

#### *Validators Dataset*

We trained a classifier for each property, e.g., one classifier is trained to classify protein sequences into different categories of subcellular localization (multiclass). To train these validator classifiers, we extracted sequences and their corresponding properties from UniProt. Specifically, we sampled the training data for the generator and the validator from the same pool. Because these models are trained in parallel, the testing data for the entire pipeline were not used to train the generator or the validators.

**Subcellular localization:** We extracted 19,225,898 and 40,000 proteins that included subcellular localization attributes in their description for the training and test sets, respectively.

**Higher taxonomic level:** Each protein in the UniProt dataset, has a taxonomy lineage attribute. The first level divides the proteins into four categories: “viruses”, “bacteria”, “archaea” and “eukaryota”. We extracted 35,632,741 and 40,000 proteins with taxonomic level attributes, for the training and test datasets, respectively.

**Enzyme:** We extracted proteins belonging to an enzyme family and proteins with catalytic activities. We also extracted the same number of proteins that show no enzymatic activity. The training data contained 51,681,180 proteins, half of which were tagged as enzymes and half of which were tagged as non-enzymes. The test data included 20,000 proteins with enzymatic / catalytic activity and 20,000 proteins without such activity.

### Supplementary Information S4: Judge Prompts

Rejecting unlikely descriptions comprises two parts. First, we wrote a Python script to verify that those proteins that the validator determined as enzymes have enzymatic activity in their description and vice versa: non-enzyme proteins do not include the term enzymatic activity in their description. Specifically, if either the string “catalytic activity” (this string is one of the fields in the UniProt database) or the string “belongs to” followed by “enzyme family” are in the description, we determine the description to be of an enzyme. Otherwise, we determined the description to be of non-enzyme. If the validator and the description are congruent, we accept that description and continue to the subsequent checks by the judge. While this enzymatic activity test was rule-based, the following checks are prompt-based.

We utilize GPT4 to implement the judge. It receives as input an English description from the generator and an English text describing the results from the validators. It uses this information to reject unlikely descriptions. Input to the judge is provided as a prompt. In fact, for each description, the judge is executed up to three times. We first provide a description and the validator prediction regarding the subcellular localization. Specifically, we provide the following prompt:

> "*You're a biology expert, and you can answer only yes or no. No explanation is needed.*
> *The protein subcellular localization is probably in one or more of the following locations: {predicticted_cell_locations}*
> *Do you think the following function is possible? please answer yes or no only.*
> *{protein_function_prediction}*"

If GPT4 returns a "yes", the description will be tested against additional validator properties. Otherwise, it is rejected. Similarly to the above test, we next ask the judge about the congruence between the description and the validator's higher taxonomy level prediction:

> "*You're a biology expert, and you can answer only yes or no. No explanation is needed.*
> *We think that the following protein belongs to the {predicted_ higher_taxonomy_level}.*
> *Do you think the following function is possible? please answer yes or no only.*
> *{protein_function_prediction}"*

Lastly, if the description is valid given the previous prompt, the judge determines whether the description is possible by both properties, i.e., the higher taxonomy level and the subcellular localization combined (we combine these two properties because the subcellular localization highly depends on the taxonomy level classification):

> "*You're a biology expert, and you can answer only yes or no. No explanation is needed.*
> *We think that the following protein belongs to the {predicted_ higher_taxonomy_level}.*
> *In addition, the protein subcellular localization is probably in one or more of the following locations: {predicticted_cell_locations}*
> *Do you think the following function is possible? please answer yes or no only.*
> *{protein_function_prediction}"*

### Supplementary Information S5: Evaluation Metrics

#### *ChrF*

Character n-gram F-score (ChrF) is an evaluation metric for strings, particularly useful in the context of text translation (Popović, 2015). It quantifies the level of substring sharing between two strings. ChrF values range from 0 (low similarity) to 1 (identical strings). ChrF captures finer nuances of the text by considering substrings (unlike word-based metrics), making it more sensitive. Formally, ChrF extracts $n$-grams (substrings of length $n$) from the reference and the hypothesis strings. Then, it calculates the precision (number of matching $n$-grams divided by the total number of $n$-grams in the hypothesis string) and the recall (number of matching $n$-grams divided by the total number of $n$-grams in reference). Next, the F-score is computed, which in this case is called ChrF:

$$ChrF = (1 + \beta^2)\frac{Precision \times Recall}{\beta^2 \times Precision + Recall}$$

We used the default values of $n$ and $\beta$ from the Huggingface library (Wolf et al., 2020): $n = 6$ and $\beta = 2$.

#### *SacreBLEU*

SacreBLEU (Post, 2018) is a metric for evaluating the quality of machine-translated text by comparing it to a reference translation. It is a more standardized and reproducible version of the Bilingual Evaluation Understudy (BLEU) score, which measures the overlap of $n$-grams (contiguous sequences of words) between two strings (Papineni et al., 2002). The metric range is between 0 to 1, with higher values indicating higher similarity. In machine translation, e.g., English to Spanish, a high-quality translation tends to have scores between 0.2 and 0.5. We compute this metric using the Huggingface library (Wolf et al., 2020).

#### *Cosine Similarity Score*

The cosine similarity score measures the similarity between two strings by computing the cosine of the angle between their vector representations (embeddings) in a multidimensional space. To compute this score, we embed the strings with our generator (embeddings from the last hidden layer). The cosine similarity score is then calculated as the dot product of these vectors divided by the product of their magnitudes. The range is from -1 to 1, where a value of -1 indicates diametrically opposed strings, 0 indicates orthogonal strings with no common terms, and 1 indicates identical strings. Cosine similarity is particularly useful in text analysis and information retrieval since the metric focuses not on the strings but on their numeric representation. Thus, this metric quantifies the distance between the meanings of strings and not the characters or substrings that make up these strings.

### Supplementary Information S6: Low-Complexity Regions

In low-complexity regions, the variability in the number of different amino acids is reduced compared to typical protein regions. In most of these regions, there is a repeated motif, such as a single amino acid or a specific pattern (Mier et al., 2019). To identify such regions, we used the PlaToLoCo webserver (Jarnot et al., 2020), which computes such regions based on five programs: SEG (Wootton & Federhen, 1993), CAST (Promponas et al., 2000), fLPS (Harrison, 2017), SIMPLE (Albà et al., 2002), and GBSC (Jarnot et al., 2020). Figure S2 provides the low complexity regions computed by

PlaToLoCo for proteins A0A8C0XGC0 and A0A1E3Q8Q4. As can be seen, all programs identified low-complexity regions in these proteins.

(a)

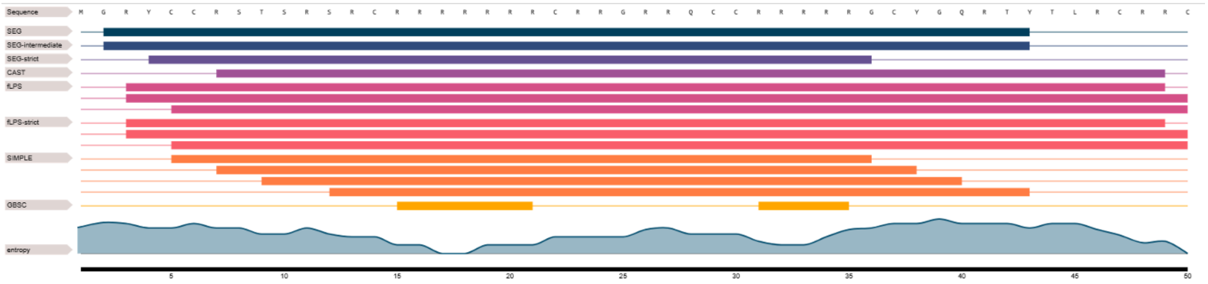


(b)

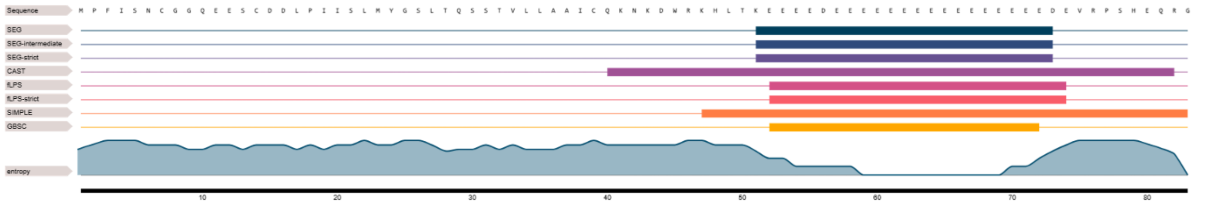


*Figure S2: Low complexity regions of proteins A0A8C0XGC0 (a) and A0A1E3Q8Q4 (b) as computed by the PlaToLoCo webserver. The first row displays the amino acids, the last row reports the entropy, and other rows correspond to the program results: a thin line corresponds to a normal region, and a bold line corresponds to a low complexity region.*

### Supplementary Information S7: Categories 1, 2 and 3 Protein Data

Supplementary Data file 1 provides the 189 Category 1 test protein identifiers and their corresponding sequence. As stated in the main text, 38 of these proteins have identical sequences in the training set. Those proteins are: P0DL03, P85622, P86130, P85660, P85735, P67789, P85743, P85683, P86630, P85534, P85658, B3A0A5, P85620, B0M3A2, P85732, B0M3B9, B0M8U4, P0C8P7, P85578, P84660, P85566, P85582, P85671, P68125, P85625, P84119, B0M8U1, P84662, P85676, C0HKK7, P85535, P84306, P0DM76, P86629, P85739, P85627, B3A0G6, and B3A0F5. These proteins are short (maximum length is 14), with an average length of 10.2 amino acids. The average length of the remaining proteins is much higher, exceeding 100 amino acids.

Supplementary Data file 2 provides the 172 Category 2 test protein identifiers, their corresponding sequences, their nearest hit (by BlastP), and the associated E-value. As stated in the main text, 21 of these proteins have identical sequences in the training set. Those proteins are: P0C7R8, P82619, P84410, P84412, P84413, P84416, P84419, P85602, A0A2Z1U82, B2Z6V3, B2Z6V7, B2Z6Y9, C9DD81, C9DD97, C9DDA7, F2VFH8, Q1P9V3, Q1P9X9, Q1P9Z1, F2VF75, and F2VFE7. As stated above, the proteins with exact hits in the training set are shorter than those without. Supplementary Data file 3 provides the 1,000 samples Category 3 test protein identifiers and their corresponding sequences, with their BlastP nearest hit and the E-value result.

### Supplementary Information S8: Detailed Performance on Category 2 and 3 Results

Table S3 provides detailed scores of ChrF, SacreBLEU, and cosine similarity and the number of exact matches (cases in which the output description perfectly matches the true one) for Category 2 (proteins with insignificant hits) and Category 3 (proteins with significant hits). For Category 2 proteins, we compared BetaDescribe's top predictions (prediction 1) and the yielded by the BlastP hit. This comparison shows a higher cosine similarity score for BetaDescribe (0.58 vs. 0.48) but lower ChrF (0.27 vs. 0.28) and SacreBLEU (0.09 vs. 0.1) scores. When testing the differences using a paired t-test, only the cosine similarity score was statistically significant (p-values: cosine similarity $< 0.0001$; ChrF $= 0.62$; SacreBLEU $= 0.84$). The hit provided by BlastP performed significantly better than BetaDescribe on all metrics when comparing the performance of Category 3 proteins.

*Table S3: Average scores of ChrF, SacreBLEU, and cosine similarity, and the count for exact matches for Categories 2 (a) and 3 (b). Predictions 1, 2, and 3 are top predictions generated by BetaDescribe.*

(a)

| | **Prediction 1 (147)** | **Prediction 2 (108)** | **Prediction 3 (87)** | **BlastP (147)** |
|---|---|---|---|---|
| **Exact Match (count)** | 2 | 3 | 0 | 5 |
| **ChrF** | $0.27 \pm 0.03$ | $0.29 \pm 0.03$ | $0.24 \pm 0.03$ | $0.28 \pm 0.03$ |
| **SacreBLEU** | $0.09 \pm 0.03$ | $0.11 \pm 0.04$ | $0.07 \pm 0.02$ | $0.1 \pm 0.03$ |
| **Cosine similarity** | $0.58 \pm 0.02$ | $0.57 \pm 0.03$ | $0.55 \pm 0.03$ | $0.48 \pm 0.03$ |

(b)

| | **Prediction 1 (910)** | **Prediction 2 (443)** | **Prediction 3 (285)** | **BlastP (910)** |
|---|---|---|---|---|
| **Exact Match (count)** | 383 | 104 | 12 | 627 |
| **ChrF** | $0.65 \pm 0.02$ | $0.52 \pm 0.03$ | $0.4 \pm 0.03$ | $0.82 \pm 0.02$ |
| **SacreBLEU** | $0.54 \pm 0.03$ | $0.37 \pm 0.04$ | $0.21 \pm 0.03$ | $0.79 \pm 0.02$ |
| **Cosine similarity** | $0.78 \pm 0.01$ | $0.7 \pm 0.02$ | $0.63 \pm 0.02$ | $0.92 \pm 0.01$ |

### Supplementary Information S9: An example in which the BlastP-based Description is More Accurate than that of BetaDescribe

In nine out of the 147 Category 2 proteins, the BlastP-based description had a higher cosine similarity score (above 0.6) than all predictions by BetaDescribe. Protein A0A3G3C7W0 exemplifies such a case. The description from UniProt is as follows:

"***FUNCTION$*** *Probable toxin that inhibits ion channels,* ***SUBCELLULAR LOCATION$*** *Secreted,* ***DOMAIN$*** *The cysteine framework is VI/VII (C-C-CC-C-C), & The presence of a 'disulfide through disulfide knot' structurally defines this protein as a knottin,* ***PTM$*** *Mostly non-hydroxylated.*"

The BlastP-based description in the training set is: "***FUNCTION$*** *Insecticidal toxin that reversibly and voltage-independendly blocks both mid-low- (M-LVA) and high-voltage-activated (HVA) calcium channels (Cav) in cockroach DUM neurons. Also causes a modest block of insect sodium channel currents (Nav). Induces potent excitatory symptoms, followed by flaccid paralysis leading to death in house crickets (By similarity),* ***SUBCELLULAR LOCATION$*** *Secreted,* ***DOMAIN$*** *The presence of a 'disulfide through disulfide knot' structurally defines this protein as a knottin,* ***MISCELLANEOUS$*** *This toxin comes from a female specimen. It is observed that propeptide sequences coming from female specimen have only limited homology with the male paralogs, but the reason is unknown,* ***SIMILARITY$*** *Belongs to the neurotoxin 08 (Shiva) family. 01 (omega toxin) subfamily.*"

The descriptions generated by BetaDescribe is: "***FUNCTION$*** *Has antibacterial activity,* ***SUBCELLULAR LOCATION$*** *Secreted,* ***SIMILARITY$*** *Belongs to the beta-defensin family.*";

"***FUNCTION$*** *Involved in gametogenesis and steroidogenesis,* ***SUBCELLULAR LOCATION$*** *Secreted,* ***SUBUNIT$*** *Heterodimer of an alpha and a beta chain,* ***SIMILARITY$*** *Belongs to the glycoprotein hormones subunit beta family.*";

"***FUNCTION$*** *Has antibacterial activity,*"

### Supplementary Information S10: Ablation Tests

#### *Effect of Pretraining on Performance*

The final training of our generator is termed Stage 3 (See main text and Supplementary Information S1). The Preliminary Stage, Stage 1, and Stage 2 are pretraining used to incorporate English and protein knowledge within our model. Here, we evaluated whether such pretraining contributed to performance. Specifically, we evaluated the performance of the model if instead of using pretraining, we initiated Stage 3 with randomly initialized weights. This model was trained on approximately 8.3 billion tokens, equivalent to 2,000 steps. The model struggled to generate any properly structured descriptions and had a failure rate of 98.5% (no function predicted in the description). Among the 24 validation proteins for which the model successfully predicted the function, eight had the exact protein sequence in the training set. In contrast, the model trained on Stages 1 and 2, using the same 2,000 steps, failed to predict only 34.3% of the test set, clearly demonstrating the importance of pretraining.

#### *The Validators Performance*

We evaluated the performance of the validators on a validation set comprising 40,000 proteins. Table S4 reports the precision error rate (1 minus precision) for the subcellular localization validator and the error rate for the higher taxonomy and enzymatic activity validators. The subcellular localization prediction is a multi-label classification task; thus, the accuracy is extremely high (0.99993). Hence, we report the precision error for that validator. As can be seen, the validators' error reaches a plateau and stabilizes around 0.011, 0.0035, and 0.027 for the subcellular localization, higher taxonomy level, and enzymatic activity predictions, respectively. This indicates that enzyme classification is more challenging than the higher taxonomy level classification. These low error rates explain the importance of including the validator as part of the pipeline: the low error rates allow the judge, relying on the validator's predictions, to reject unlikely generated descriptions reliably.

*Table S4: Performance of the different validators as a function of the number of training steps. For subcellular localization, we report the precision error rate (one minus precision). For the two other categories, we report error rates (one minus accuracy). Values are calculated using a validation set of 40,000 proteins.*

| Steps | 120K | 240K | 360K | 480K | 600K | 720K | 840K | 960K | 1,080K | 1,200K |
|---|---|---|---|---|---|---|---|---|---|---|
| **Subcellular localization** | 0.023 | 0.02 | 0.017 | 0.015 | 0.014 | 0.014 | 0.013 | 0.012 | 0.012 | 0.012 |
| **Higher taxonomy level** | 0.0133 | 0.0091 | 0.0077 | 0.0063 | 0.0057 | 0.005 | 0.0046 | 0.004 | 0.0037 | 0.0035 |
| **Enzymatic activity** | 0.049 | 0.042 | 0.035 | 0.034 | 0.033 | 0.031 | 0.034 | 0.028 | 0.029 | 0.027 |

#### *The Judge Performance*

To evaluate the judge, we manually curated the following dataset. First, we randomly sampled 200 descriptions from the training dataset, each with a subcellular localization. Each description is matched with a pair of properties mimicking the validator output. An example of such a pair of properties is "viruses" and "endoplasmic reticulum". We manually modified the properties of some of these pairs so that 100 descriptions are matched with "true" properties, i.e., the properties match the description provided by UniProt. For the remaining 100 descriptions, we modified at least one of the properties so that the validator and the descriptions are incongruent. We expect the judge to correctly accept the first 100 samples and reject the others. For example, consider the following description:

***FUNCTION$*** *Inhibits post-transcriptional processing of cellular pre-mRNA, by binding and inhibiting two cellular proteins that are required for the 3'-end processing of cellular pre-mRNAs: the 30 kDa cleavage and polyadenylation specificity factor/CPSF4 and the poly(A)-binding protein 2/PABPN1. In turn, unprocessed 3' end pre-mRNAs accumulate in the host nucleus and are no longer exported to the cytoplasm. […],* ***SUBUNIT$*** *Homodimer. Interacts with host TRIM25 (via coiled coil); this interaction specifically inhibits TRIM25 multimerization and TRIM25-mediated RIGI CARD ubiquitination. Interacts with human EIF2AK2/PKR, CPSF4, IVNS1ABP and PABPN1,* ***SUBCELLULAR LOCATION$*** *Host nucleus,* ***DOMAIN$*** *The dsRNA-binding region is required for suppression of RNA silencing,* ***SIMILARITY$*** *Belongs to the influenza A viruses NS1 family.*

With the real properties of "viruses" and the subcellular localization of the "host nucleus", we expect the judge to accept the description correctly. We expect the judge to reject the description with false properties, such as a higher taxonomy level of “eukaryote” and a subcellular location of “endoplasmic reticulum”. Out of the 100 descriptions with the properties from the training set, the judge (GPT4) correctly classified 81. Out of the 100 descriptions with erroneous (altered) properties, the judge correctly rejected 87. Thus, the accuracy of GPT4 is 0.84, with a precision of 0.86, and an F1 score of 0.84.

### Supplementary Information S11: Public LLMs Predictions of the Function of Proteins Given a Protein Sequence as Input

We tested how public LLMs perform compared to BetaDescribe, i.e., how well they can describe the function of a given protein when provided its sequence as input. We used the public versions as of October 2024 of the following models: GPT4 (OpenAI, 2023), Gemini 1.5 Flash (Gemini Team et al., 2024), and Claude 3.5 Sonnet. The following prompt was used for this experiment:

“*What is the function of the protein with the following sequence:*

*{amino_acid_chain}*

*Use the following output format.*

*Example 1 of the output format:*

*FUNCTION$ Catalyzes the reduction of fatty acyl-CoA to fatty alcohols, CATALYTIC ACTIVITY$ a long-chain fatty acyl-CoA + 2 H(+) + 2 NADPH = a long-chain primary fatty alcohol + CoA + 2 NADP(+), SIMILARITY$ Belongs to the fatty acyl-CoA reductase family.*

*Example 2 of the output format:*

*FUNCTION$ Nuclease required for the repair of DNA interstrand cross-links (ICL). Acts as a 5'-3' exonuclease that anchors at a cut end of DNA and cleaves DNA successively at every third nucleotide, allowing to excise an ICL from one strand through flanking incisions, CATALYTIC ACTIVITY$ Hydrolytically removes 5'-nucleotides successively from the 3'-hydroxy termini of 3'-hydroxy-terminated oligonucleotides., COFACTOR$ Mg(2+), SUBCELLULAR LOCATION$ Nucleus, SIMILARITY$ Belongs to the FAN1 family.*

*Example 3 of the output format:*

*FUNCTION$ Directs RNA polymerase II nuclear import, SUBCELLULAR LOCATION$ Cytoplasm, SIMILARITY$ Belongs to the IWR1/SLC7A6OS family.*”

Supplementary Data file 4 provides the resulting prediction for each protein. We note that answers may contain unrelated information (such as suggestions for using bioinformatics tools). Thus, we manually processed their answers to extract only the meaningful descriptions for the evaluation metrics.

### Supplementary Information S12: Descriptions Provided by BetaDescribe and BlastP Search for Unknown Proteins

Table S5 provides the predictions generated by BetaDescribe as well as the BlastP-based prediction for four proteins for which the function was not experimentally validated at the time of BetaDescribe analyses: SnRV-Env, TiLV Protein 1, TiLV Protein 2, and TiLV Protein 3.

*Table S5: Detailed descriptions provided by BetaDescribe (Predictions 1, 2, and 3) and the BlastP-based prediction. Panels (a), (b), (c), and (d) correspond to SnRV-Env protein, TiLV Protein 1, TiLV Protein 2, and TiLV Protein 3, respectively.*

(a)

| | |
|---|---|
| BetaDescribe (Prediction 1) | ***FUNCTION$*** *The leader peptide is a component of released, infectious virions and is required for particle budding, & The transmembrane protein (TM) acts as a class I viral fusion protein. Under the current model, the protein has at least 3 conformational states: pre-fusion native state, pre-hairpin intermediate state, and post-fusion hairpin state. During viral and target cell membrane fusion, the coiled coil regions (heptad repeats) assume a trimer-of-hairpins structure, positioning the fusion peptide in close proximity to the C-terminal region of the ectodomain. The formation of this structure appears to drive apposition and subsequent fusion of viral and target cell membranes. Membranes fusion leads to delivery of the nucleocapsid into the cytoplasm,* ***SUBCELLULAR LOCATION$*** *Endoplasmic reticulum membrane.* |
| BetaDescribe (Prediction 2) | ***FUNCTION$*** *Acts as a ligand for KLRK1,* ***SUBCELLULAR LOCATION$*** *Cell membrane,* ***SIMILARITY$*** *Belongs to the NKG2D ligand family.* |
| BlastP (protein A0A0R1ZYJ7, E-value of 0.5) | ***FUNCTION$*** *Responsible for synthesis of pseudouridine from uracil,* ***CATALYTIC ACTIVITY$*** *a uridine in RNA = a pseudouridine in RNA,* ***SIMILARITY$*** *Belongs to the pseudouridine synthase RluA family.* |

(b)

| | |
|---|---|
| BetaDescribe (Prediction 1) | ***FUNCTION$*** *Probably involved in the RNA silencing pathway and required for the generation of small interfering RNAs (siRNAs),* ***CATALYTIC ACTIVITY$*** *a ribonucleoside 5'-triphosphate + RNA(n) = diphosphate + RNA(n+1),* ***SIMILARITY$*** *Belongs to the RdRP family.* |
| BetaDescribe (Prediction 2) | ***FUNCTION$*** *Hydrolase that can remove 'Lys-48'-linked conjugated ubiquitin from proteins,* ***CATALYTIC ACTIVITY$*** *Thiol-dependent hydrolysis of ester, thioester, amide, peptide and isopeptide bonds formed by the C-terminal Gly of ubiquitin (a 76-residue protein attached to proteins as an intracellular targeting signal).,* ***SIMILARITY$*** *Belongs to the MINDY deubiquitinase family. FAM188 subfamily.* |
| BetaDescribe (Prediction 3) | ***FUNCTION$*** *The proteasome is a multicatalytic proteinase complex which is characterized by its ability to cleave peptides with Arg, Phe, Tyr, Leu, and Glu adjacent to the leaving group at neutral or slightly basic pH. The proteasome has an ATP-dependent proteolytic activity. This subunit is involved in antigen processing to generate class I binding peptides,* ***CATALYTIC ACTIVITY$*** *Cleavage of peptide bonds with very broad specificity.,* ***SUBCELLULAR LOCATION$*** *Nucleus.* |
| BlastP (protein O41353, E-value of 0.000871) | ***FUNCTION$*** *RNA-dependent RNA polymerase which is responsible for replication and transcription of virus RNA segments. The transcription of viral mRNAs occurs by a unique mechanism called cap-snatching. 5' methylated caps of cellular mRNAs are cleaved after 10-13 nucleotides by PA. In turn, these short capped RNAs are used as primers by PB1 for transcription of viral mRNAs. During virus replication, PB1 initiates RNA synthesis and copy vRNA into complementary RNA (cRNA) which in turn serves as a template for the production of more vRNAs,* ***CATALYTIC ACTIVITY$*** *a ribonucleoside 5'-triphosphate + RNA(n) = diphosphate + RNA(n+1),* ***SUBUNIT$*** *RNA polymerase is composed of three subunits: PA, PB1 and PB2,* ***SIMILARITY$*** *Belongs to the influenza viruses polymerase PB1 family.* |

(c)

| | |
|---|---|
| BetaDescribe (Prediction 1) | ***FUNCTION$*** *DNA-dependent RNA polymerase catalyzes the transcription of DNA into RNA using the four ribonucleoside triphosphates as substrates,* ***CATALYTIC ACTIVITY$*** *a ribonucleoside 5'-triphosphate + RNA(n) = diphosphate + RNA(n+1),* ***SIMILARITY$*** *Belongs to the RNA polymerase beta' chain family.* |

| | |
|---|---|
| BlastP (protein A0A357NC89, E-value of 3.4) | ***FUNCTION$*** *Synthesizes alpha-1,4-glucan chains using ADP-glucose,* ***CATALYTIC ACTIVITY$*** *[(1->4)-alpha-D-glucosyl](n) + ADP-alpha-D-glucose = [(1->4)-alpha-D-glucosyl](n+1) + ADP + H(+),* ***PATHWAY$*** *Glycan biosynthesis; glycogen biosynthesis,* ***SIMILARITY$*** *Belongs to the glycosyltransferase 1 family. Bacterial/plant glycogen synthase subfamily.* |

(d)

| | |
|---|---|
| BetaDescribe (Prediction 1) | ***FUNCTION$*** *DNA-dependent RNA polymerase catalyzes the transcription of DNA into RNA using the four ribonucleoside triphosphates as substrates. Specific core component of RNA polymerase III which synthesizes small RNAs, such as 5S rRNA and tRNAs,* ***SUBCELLULAR LOCATION$*** *Nucleus,* ***SUBUNIT$*** *Component of the RNA polymerase III (Pol III) complex consisting of 17 subunits,* ***SIMILARITY$*** *Belongs to the eukaryotic RPC3/POLR3C RNA polymerase subunit family.* |
| BetaDescribe (Prediction 2) | ***FUNCTION$*** *Interacts with EME1 to form a DNA structure-specific endonuclease with substrate preference for branched DNA structures with a 5'-end at the branch nick. Typical substrates include 3'-flap structures, D-loops, replication forks and nicked Holliday junctions. May be required in mitosis for the processing of stalled or collapsed replication fork intermediates. May be required in meiosis for the repair of meiosis-specific double strand breaks subsequent to single-end invasion (SEI),* ***SUBCELLULAR LOCATION$*** *Nucleus,* ***COFACTOR$*** *Mg(2+),* ***SUBUNIT$*** *Interacts with EME1,* ***SIMILARITY$*** *Belongs to the XPF family.* |
| BetaDescribe (Prediction 3) | ***FUNCTION$*** *Decapping enzyme for NAD-capped RNAs: specifically hydrolyzes the nicotinamide adenine dinucleotide (NAD) cap from a subset of RNAs by removing the entire NAD moiety from the 5'-end of an NAD-capped RNA,* ***SUBCELLULAR LOCATION$*** *Nucleus,* ***COFACTOR$*** *a divalent metal cation,* ***SIMILARITY$*** *Belongs to the DXO/Dom3Z family.* |
| BlastP (protein A0A1F6CNS5, E-value of 1.8) | ***FUNCTION$*** *Associates with the EF-Tu.GDP complex and induces the exchange of GDP to GTP. It remains bound to the aminoacyl-tRNA.EF-Tu.GTP complex up to the GTP hydrolysis stage on the ribosome,* ***SUBCELLULAR LOCATION$*** *Cytoplasm,* ***SIMILARITY$*** *Belongs to the EF-Ts family.* |

### Supplementary Information S13: Quantifying the Functionally Importance of Protein Regions

We employed *in-silico* alanine scanning mutagenesis, using a sliding-window approach, with a window size of ten amino acids and a shift of one amino acid, i.e., overlapping windows. To quantify the importance of substituting the residues in a specific sequence window to alanine, we used the negative log-likelihood metric, which provides a fit between a sequence and a description (see below). Thus, for each amino acid, there are ten values (except the ones in the edges of the protein sequence) of negative-log likelihood scores (one for each participation in a window). We averaged the negative-log likelihood for each amino acid and normalized the scores via a power transformation (Yeo & Johnson, 2000), which averaged the scores to zero. Next, we subtracted the normalized value of the wild-type protein sequence, such that the average score of the wild-type sequence becomes zero. We note that we did not train a model to predict the impact of mutagenesis experiments. Instead, we used an unsupervised approach that relies on the previous training of the BetaDescribe generator to analyze the mutated sequences.

#### *Negative Log-Likelihood*

In NLP, negative log-likelihood measures how well a model predicts text. It is commonly used to evaluate language models by determining how "surprised" the model is by specific tokens. A lower negative log-likelihood indicates that the model is more confident in its predictions, assigning higher probabilities to the actual next tokens in the sequence. Models are trained to minimize this value and thus improve language understanding and generation accuracy. For the following $n$ tokens: $w_1, w_2, \dots, w_n$, the negative log-likelihood is defined as:

$$negative\ log\ likelihood = -\frac{1}{n}\sum_{i=1}^{n}\log(p(w_i))$$

Where $p(w_i)$ is the probability of generating the token $w_i$ given the previous tokens: $w_1 \dots w_{i-1}$.

#### *Additional Example for Identifying Functionally Important Protein Regions*

RecA of *Escherichia coli* (P0A7G6), the founding member of the bacterial RecA protein family, is essential for initiating DNA break repair, activating the SOS response, enabling translation synthesis, and promoting the spread of antibiotic resistance genes. For its various roles, RecA has a few functional regions: N-terminal domain, "Make ATP Work" (MAW) motif, A site, B site, and DNA binding sites (McGrew & Knight, 2003). Previous studies showed that mutating these regions may lead to severe defects in RecA function (Lee & Wang, 2009; Leite et al., 2019; McGrew & Knight, 2003). As reported in Figure S3, BetaDescribe indicated that mutations in some of the functional sites, such as the N-terminal domain, MAW motif, and A site, increase the negative log-likelihood of the description. In other words, BetaDescribe was mainly influenced by mutations in certain functional regions and significantly less so by mutations in the non-functional areas. However, not all functional regions were identified as important. The B site and some binding sites were not classified as important. These analyses suggest that the BetaDescribe model can be used in an unsupervised approach to identify regions of importance with relatively high accuracy.

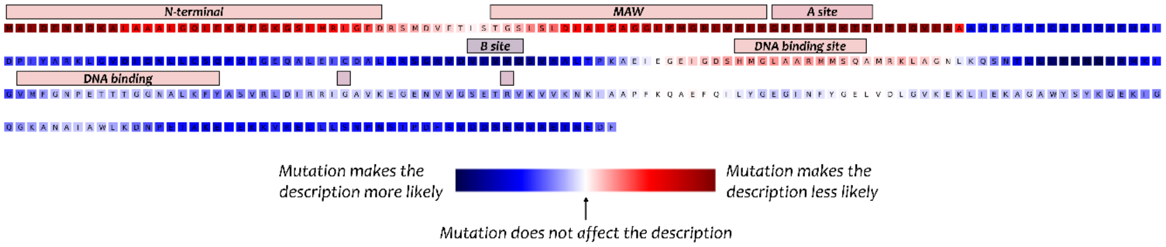


*Figure S3: Functionally Important regions within the RecA protein as identified by BetaDescribe. The importance of each region is computed by a window-sliding, in-silico mutagenesis approach. The negative log-likelihood of each residue is evaluated to quantify functional importance. The values are normalized such that zero represents the average conservation score of the wild-type protein. The annotation of each functionally important region is stated above its position, except for two binding sites (G229 and R243).*
